# Joint eQTL mapping and Inference of Gene Regulatory Network Improves Power of Detecting both *cis*- and *trans*-eQTLs

**DOI:** 10.1101/2020.04.23.058735

**Authors:** Xin Zhou, Xiaodong Cai

## Abstract

**Motivation:** Genetic variations of expression quantitative trait loci (eQTLs) play a critical role in influencing complex traits and diseases development. Two main factors that affect the statistical power of detecting eQTLs are: 1) relatively small size of samples available, and 2) heavy burden of multiple testing due to a very large number of variants to be tested. The later issue is particularly severe when one tries to identify *trans*-eQTLs that are far away from the genes they influence. If one can exploit co-expressed genes jointly in eQTL-mapping, effective sample size can be increased. Furthermore, using the structure of the gene regulatory network (GRN) may help to identify *trans*-eQTLs without increasing multiple testing burden.

**Results:** In this paper, we employ the structure equation model (SEM) to model both GRN and effect of eQTLs on gene expression, and then develop a novel algorithm, named sparse SEM, for eQTL mapping (SSEMQ) to conduct joint eQTL mapping and GRN inference. The SEM can exploit co-expressed genes jointly in eQTL mapping and also use GRN to determine *trans*-eQTLs. Computer simulations demonstrate that our SSEMQ significantly outperforms eight existing eQTL mapping methods. SSEMQ is further employed to analyze a real dataset of human breast tissues, yielding a number of *cis*- and *trans*-eQTLs.

**Availability:** R package ssemQr is available on https://github.com/Ivis4ml/ssemQr.git.

## Introduction

A long-standing problem in genetics is to understand how genetic variation affects phenotypic variation. As central dogma of molecular biology indicates, the information encoded in DNA sequences flows through message RNA to protein. Consequently, DNA sequence polymorphisms may influence final phenotypes through intermediate, molecular phenotypes such as gene expression levels and protein structures. Understanding the genetic architecture of gene expression is therefore of paramount importance to the understanding of the molecular origins of complex traits and diseases (Albert and Kruglyak, 2015). eQTL mapping has been used to identify genetic variants that affect gene expression. Genome-wide eQTL mapping needs to test a very large number of variants, which results in a heavy burden of multiple testing and low statistical power. For this reason, many eQTL mapping studies are limited to identify *cis*-eQTLs that are located close, typically within 1 megabases (Mb), to the genes (*cis*-eGenes) that they influence. This significantly reduces the number of variants to be tested, which, together with the fact that *cis*-eQTLs typically have large effect sizes, improve the power of detecting eQTLs. However, *cis*-eQTLs explain a relatively small fraction of total gene expression heritability (Grundberg *et al*., 2012), and a substantial proportion of gene expression heritability can be due to *trans*-eQTLs that are far away from the genes (*trans*-eGenes) they influence, often on a different chromosome (Grundberg *et al*., 2012). Therefore, to better understand the genetic base of gene expression, it is critical to identify *trans*-eQTLs.

Identification of *trans*-eQTLs is however challenging due to the heavy multiple testing burden and the fact that the effect sizes of *trans*-eQTLs are typically small. A recent study shows that a large number of eQTLs regulate gene expression in both a *cis* and *trans* manner, and that about one third *trans* regulation is mediated by the expression of *cis*-eGenes (Yao *et al*., 2017). Several methods have been developed recently to identify pairs of *trans*-eQTL and *trans*-eGenes that are mediated by *cis*-eGenes, based on a mediation test (Shan *et al*., 2019; Yang *et al*., 2017, 2019) or expression levels of possible mediator genes predicted from their *cis*-eQTLs (Liu *et al*., 2018; Wheeler *et al*., 2019). However, a mediation test relies on *trans*-eQTL and *trans*-eGenes pairs already detected by an existing method which may still suffer from a heavy multiple testing burden, and the power of the methods relying on predicted gene expression levels may be affected negatively by prediction errors in gene expression levels.

If the structure of the gene regulatory network (GRN) of the organism under consideration is known or can be inferred from gene expression data available, once a pair of *cis*-eQTL and *cis*-eGene is identified, the *trans*-eGenes mediated by the *cis*-eGene can be determined through the GRN. In this paper, we employ the structure equation model (SEM) to model both GRN and effect of eQTLs on gene expression, and then develop a novel algorithm, named sparse SEM for eQTL mapping (SSEMQ), to conduct joint eQTL mapping and GRN inference. The algorithm outputs *cis*-eQTLs and the GRN structure, which are further used to determine *trans*-eGenes mediated by *cis*-eGenes. As SSEMQ exploits correlated expression of multiple genes jointly, it effectively increases the sample size, thereby improving the power of detecting eQTLs. As *trans*-eQTLs and *trans*-eGenes are determined by the GRN, this approaches doe not increase multiple testing burden, which can improve the power of detecting *trans*-eQTLs. Indeed, computer simulations demonstrate that our SSEMQ significantly outperforms eight existing eQTL mapping methods. The SSEMQ is further employed to analyze a genotype and gene expression dataset of human breast tissues from the Genotype-Tissue Expression (GTEx) Project (Lonsdale *et al*., 2013), yielding a number of *cis*- and *trans*-eQTLs. Of note, SEM was employed recently to conduct genome-wide association studies (GWAS) (Momen *et al*., 2018; Wang *et al*., 2019).

## Methods

### 2.1 SEM for eQTL mapping

We will consider the problem of identifying *cis*-eQTLs that are located within *L* bps of a gene (*cis*-eGene), and possibly act as a *trans*-eQTLs to affect the expression of other remote genes (*trans*-eGenes) through the mediation of the *cis*-eGene. The distance *L* depends on the organism under consideration; it is typically 1 M bps for *Homo sapiens* (Kirsten *et al*., 2015), or 515 bps for yeast *S*.*cerevisiae* (Sunnerhagen and Piskur, 2006). Figure 1 depicts such an example, where the SNP is a *cis*-eQTL of the *cis*-eGene that regulates the expression of two remote genes *trans*-eGene_1_ and *trans*-eGene_2_ directly, and *trans*-eGene_1_ also regulates the expression of another gene, *trans*-eGene_3_, through GRN. Therefore, the SNP is not only a *cis*-eQTL of the *cis*-eGene, but also a *trans*-eQTL of *trans*-eGenes 1, 2, and 3. Of note, an eQTL can influence a *trans*-eGene through other mechanisms, but they will not be considered in this paper.

**Figure 1.**
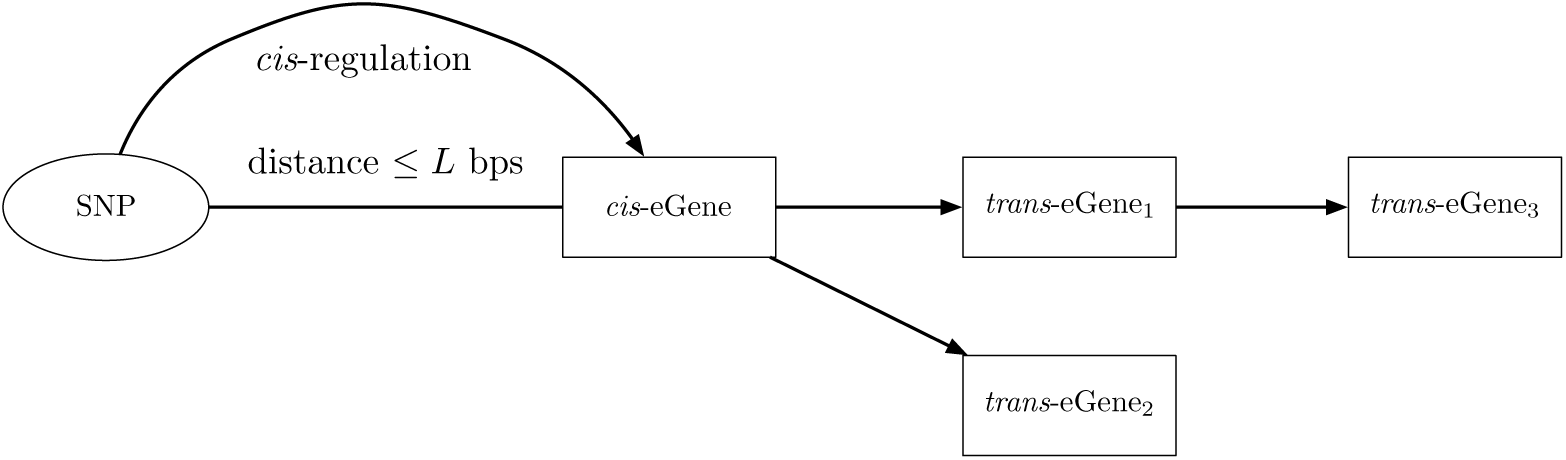
Regulatory Mechanisms of eQTLs considered in this paper. Genetic variants can affect genes expressions through the following mechanisms: (1) SNP affects local gene expression (*cis*-eGene); (2) SNP affects remote genes (*trans*-eGene) expressions by mediation of a *cis*-eGene.

Suppose that the expression levels of *K* genes in *N* individuals have been collected using e.g. micro-array or RNA-Seq. Let **y**_*i*_ = [*y*_*i*1_, *y*_*i*2_, …, *y*_*iK*_]^*T*^ denote the expression levels of *K* genes in the *i*th individual, where *i* = 1, 2, …, *N*. Suppose all SNPs in the genomes of *N* individuals have been obtained. Denote the genotypes of *n*_*k*_ SNPs within a distance of *L* bps to gene *k* in the *i*th individual as 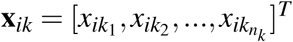, where *x*_*ikj*_ takes value from {0, 1} for haploid organisms or from {0, 1, 2} for diploid organisms. Define a *J* × 1 vector 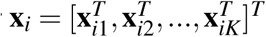, where 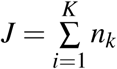. Since the expression level of a particular gene can be regulated by other genes and affected by eQTLs, we use the following SEM to model gene expression levels:

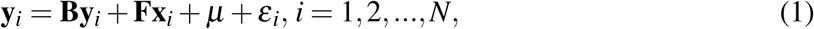

where **B** *∈* ℝ^*K*×*K*^ reflects the regulatory effects among genes; **F** *∈* ℝ^*K*×*J*^ captures the effect of possible *cis*-eQTLs on gene expression levels; *µ* ∈ℝ^*K*×1^ accounts for the model bias and *ε*_*i*_ ∈ℝ^*K*×1^ is the residual error, which is assumed as an Gaussian random vector with zero mean and covariance matrix *σ* ^2^**I**, with **I** being the *K* × *K* identity matrix. Matrix **B** characterizes the structure of the GRN, and it is assumed that there is no self-loops presented in GRN (Logsdon and Mezey, 2010), which implies that diagonal entries of **B** are zero. Define a set *S*_*i*_ = {*m*_*i*_ + 1, *m*_*i*_ + 2, …, *m*_*i*_ + *n*_*i*_} that contains the indices of SNPs within a distance of *L* bps to the *i*th gene, where *m*_*i*_ = 0 if *i* = 1 or 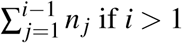. Then *F*_*i j*_ = 0 if *j* ∉ *S*_*i*_, whereas **F**_*i j*_, *i* = 1, …, *K, j S*_*i*_ are model parameters to be estimated.

Of note, the same SEM was employed to infer GRNs in (Cai *et al*., 2013; Logsdon and Mezey, 2010), where it is assumed that eQTLs of genes are known, and thus **x**_*i*_ contains the genotypes of known eQTLs. Here, eQTLs are unknown, therefore, **x**_*i*_ contains the genotypes of all SNPs within *L* bps of their corresponding genes. The goal of (Cai *et al*., 2013; Logsdon and Mezey, 2010) is to estimate **B**, which gives the structure of the GRN. The main purpose here is eQTL-mapping, which will be accomplished through joint estimation of **B** and **F**. After **B** and **F** are estimated, the nonzero entries of **F** will yield *cis*-eQTLs and their corresponding *cis*-eGenes; Matrix **B** will tells gene regulated by *cis*-eGenes, as illustrated in Figure 1, which in turn give *trans*-eGenes mediated by *cis*-eQTLs. Equivalently, we can form a matrix **G** = (**I**−**B**)^−1^**F**, and nonzero entries of **G** will give all pairs of eQTLs and eGenes.

### 2.2 Sparsity-aware inference of SEM

Let **Y** = [**y**_1_, **y**_2_, …, **y**_*N*_] be the gene expression data of *N* individuals, and **X** = [**x**_1_, **x**_2_, …, **x**_*N*_] ∈ ℝ^*J*×*N*^ be the genotype data. The SEM in (1) can be compactly written as **Y** = **BY** + **FX** + *µ***1**^*T*^ + **E**, where **1** is a column vector whose elements are all 1, and **E** = [*ε*_1_, …, *ε*_*N*_] ∈ ℝ^*K*×*N*^. Then, the negative log-likelihood function can be written as

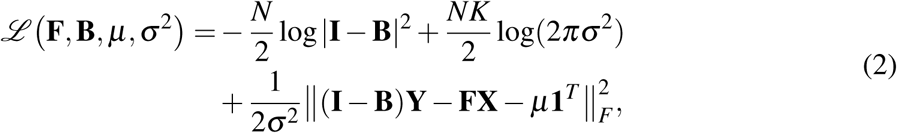

where | · | denotes matrix determinant and ‖.· ‖_*F*_ denotes the Frobenius norm. It is not difficult to show that minimizing (2) with respect to (w.r.t.) *µ* yields 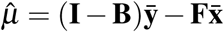, where 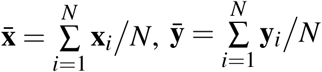.

Since the expression level of a gene is regulated by a small number of other genes (Tegner *et al*., 2003) and influenced by a small number of *cis*-eQTLs (Brem and Kruglyak, 2005; Fagny *et al*., 2017), matrices **B** and **F** are expected to be sparse, meaning that most of their entries are zero. Therefore, taking into account the sparsity property of **B** and **F**, we estimate **B** and **F** by solving the following *𝓁*_1_-penalized maximum likelihood estimation problem:

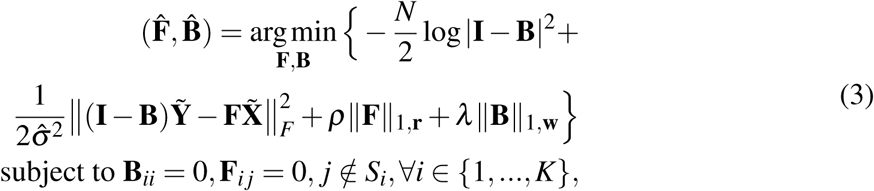

where 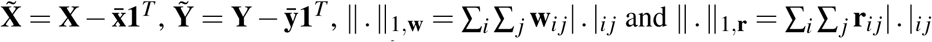 represent the weighted *𝓁*_1_-norm of a given matrix; *λ, ρ* are two nonnegative parameters. Note that the total number of unknown coefficients in **B** and **F** is *p* = *K*(*K* −1) + *KJ*, which can be much larger than the number of data sample *N*. The high dimensional optimization problem (3) is challenging, but our SSEMQ algorithm to be presented later can handle the case where *p ≫ N*. The weights **w** and **r** introduced in the penalty terms are suggested by the adaptive lasso (Zou, 2006) to improve the consistency and robustness of the estimates (Cai *et al*., 2013; Zhou and Cai, 2020); they are chosen to be 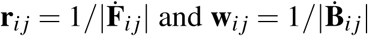, respectively, where 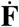 and 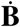 are preliminary estimates of **F** and **B**, respectively, obtained from the following ridge regression:

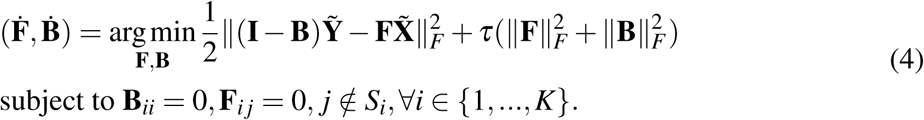

The estimate 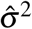 in (3) can be obtained as 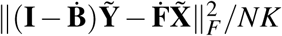.

The major difference between optimization problem (3) here and the optimization problem formulated for GRN inference in (Cai *et al*., 2013) is that the **F** matrix here contains many more columns, and a regularization term *ρ* ‖**F**‖_1,**r**_ is added to the objective function. This term is necessary for eQTL mapping, because **x**_*i*_ in (1) contains all SNPs within *L* bps of gene *i*, some of which may be eQTLs. While it is possible to modify the SML algorithm in (Cai *et al*., 2013) to solve (3), the SML algorithm is very slow to handle this problem, because it employs an element-wise coordinate descent method. In this paper, we will develop a new and much more efficient algorithm using a block coordinate descent approach.

### 2.3 Closed-form solution of Ridge regression

We first solve the ridge regression problem (4) row by row separately to find initial estimates of **F** and **B**, and weights **r** and **w** for the optimization problem in (3). Let **F**_*i*,._, **B**_*i*,._ and **Y**_*i*,._ be the *i*-th row of **F, B** and **Y**, respectively; further denote 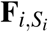 as the 1 × *n*_*i*_ vector whose elements are **F**_*i, j*_, *j ∈ S*_*i*_, and **B**_*i*,−*i*_ as the 1 × (*K* −1) vector formed by removing the *i*th element from **B**_*i*,._. Then, the ridge regression problem (4) can be decomposed into *K* separate problems as follow:

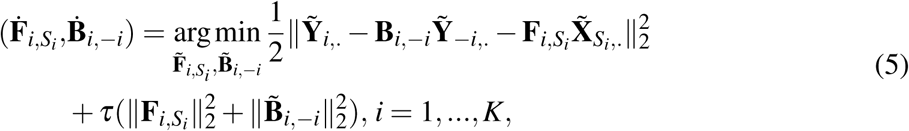

where 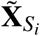,. is the submatrix obtained by deleting rows of 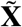 whose indices are not in *S*_*i*_, and 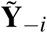,. is the matrix formed by removing the *i*th row of 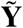. Minimizing the objective function in (5) w.r.t. 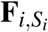 yields 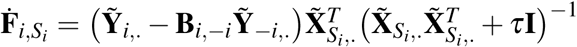. Then, substituting 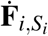 into (5) and minimizing w.r.t. **B**_*i*,−*i*_ gives 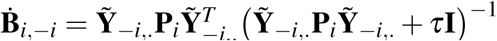, where 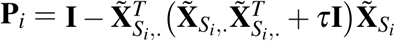. After 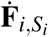 and 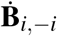 are estimated, 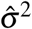 is calculated as described in section 2.3. The hyperparameter *τ* in ridge regression (4) and (5) is determined by *k*-fold cross-validation, with *k* typically being 5 or 10.

### 2.4 SSEMQ algorithm

In this section, we will develop the SSEMQ algorithm to solve the optimization problem in (3) initialized with 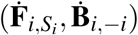 from (5). The problem is non-convex due to the log-determinant term 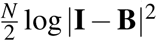, and non-smooth due to the *𝓁*_1_-norm terms of **F** and **B**. Recently, a proximal alternating linearized minimization (PALM) (Bolte *et al*., 2014) method was proposed to solve general non-convex and non-smooth problems. We will apply the PALM approach to develop our SSEMQ algorithm.

Without loss of generality, we define the proximal operator associated with a proper and lower-semi-continuous function 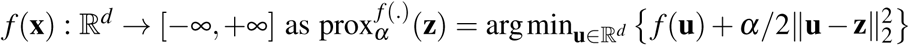, where *α* ∈ (0, +∞) and **z** ∈ℝ^*d*^ are given. If *f* (**x) =** *λ* ‖**x** ‖_1_, where *λ* > 0, then the lasso proximal operator (Parikh *et al*., 2014) can be written as follows,

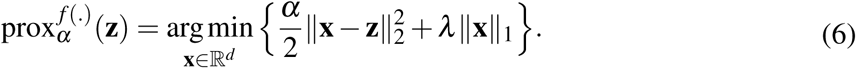

Let **x**(*λ*) denote the solution of (6) for a given *λ*, and *x*_*i*_(*λ*) denote the *i*th element of **x**(*λ*). Then, *x*_*i*_(*λ*) is given by the soft-thresholding operator as follows (Parikh *et al*., 2014):

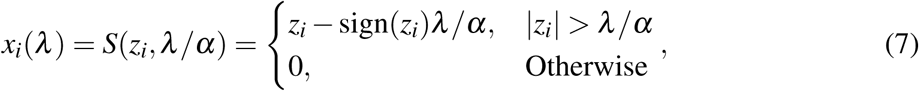

where *z*_*i*_ represents the *i*th element of **z**. For convenience, the solution of (6) can be written as 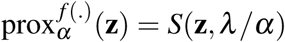, where the soft-thresholding operator is applied to **z** in an element-wise manner.

Define **B**_.,*i*_ as the *i*th column of **B, B**_−*i,i*_ as the vector formed by removing the *i*th element of **B**_.,*i*_. Let **r**_*i*,._ be the *i*th row of **r, w**_.,*i*_ be the *i*th column of **w**, and **w**_−*i,i*_ be the vector formed by removing the *i*th element of **w**_.,*i*_. To solve the optimization problem in (3) with PALM, we rewrite the objective function as

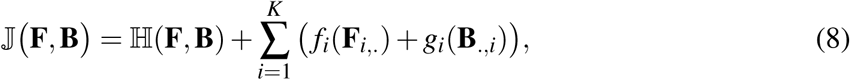

where 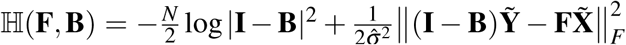, and 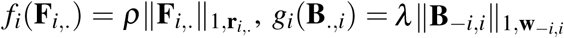.

Using the inertial version of the PALM approach, our SSEMQ algorithm efficiently minimizes 𝕁 (**F, B)** via the block coordinate descent (BCD) method in an iterative fashion. Specifically, in each cycle of the iteration, 𝕁 (**F, B)** is minimized successively w.r.t. a selected **B**_.,*i*_ or **F**_*i*,._, *i* = 1, …, *K*, while all other blocks of variables in (**F, B**) are fixed. First, let us consider updating **B**_.,*i*_ in the (*t* + 1)th iteration with other variables in **F** and **B** being fixed. Let **B**(*t*) denote the estimate of **B** in the *t*th iteration, then **B**(*t* + 1)_−*i,i*_ can be updated by lasso proximal operator defined in (6) as follows:

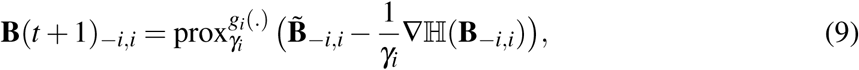

where 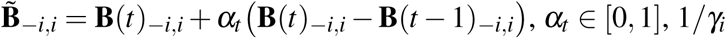 is the step-size for block **B**_−*i,i*_ that will be given later, and ∇ℍ(**B**_−*i,i*_) is the partial derivative of ℍ w.r.t. **B**_−*i,i*_ at given **B** and **F**. We can compute **B**(*t* + 1)_−*i,i*_ in (9) from (7), if we know ∇ℍ(**B**_−*i,i*_) and *γ*_*i*_, which will be derived in the following.

Let us reorganize ℍ(**F, B**) in (8) as a function of **B**_−*i,i*_, ℍ(**B**_−*i,i*_), when other var(iables in **B** and **F** are fixed. Define 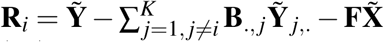, and note that log |**I** − **B**| = log **(c***i,i* − **c***i*,−*i***B**−*i,i*), where **c** *i,i* is the (*i, i*) th co-factor of **I** − **B**, and **c** *i*,−*i* contains co-factors of **I** − **B** corresponding to **B**_−*i,i*_. Then, ℍ(**B**_−*i,i*_) is defined as 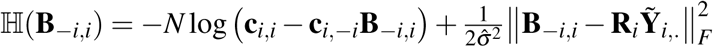, and ∇ℍ(**B**_−*i,i*_) can be written as 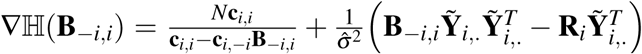. In the Supplementary text, we prove that ∇ℍ(**B**_−*i,i*_) is Lipschitz continuous. More specifically, ∇ℍ(**B**_−*i,i*_) satisfies the property that ‖∇ℍ(**x**) − ∇ℍ(**y**)‖*≤ L*_*i*_(**F, B**_.,−*i*_) ‖**x** − **y**‖, where the Lipschitz constant *L*_*i*_(**F, B**_.,−*i*_) depends on fixed variables in **F** and **B**_.,−*i*_, and is derived in the Supplementary text as

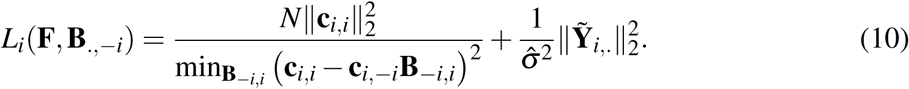

Here, the value of min**B**_−*i,i*_ (**c**_*i,i*_ − **c**_*i,−i*_**B**_−*i,i*_)^2^ can be obtained by solving the optimization problem shown in (S10) in supplementary text. The step size in (9) is then chosen as *γ*_*i*_ = *L*_*i*_(**F, B**_.,−*i*_).

Now, we consider updating the *i*th row of **F** in the (*t* + 1)th iteration with other variables being fixed. Since **F** is inside the 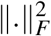 term, **F**_*i*,._ can be updated by using the standard lasso proximal operator as follows,

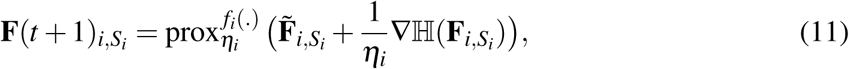

where 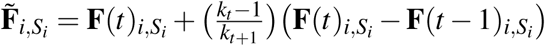, with 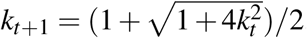, as suggested in (Beck and Teboulle, 2009), and ∇ℍ(**F**_*i,S_i_*_) is the partial derivative of ℍ w.r.t. **F**_*i,S_i_*_ with other variables fixed, and it can be written as 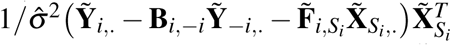,.. The step-size *η*_*i*_ can be found as 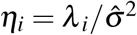, with *λ* _*i*_ being the largest eigenvalue of matrix 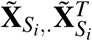.

The SSEMQ algorithm is summarized in Algorithm 1, and the convergence criterion is defined as 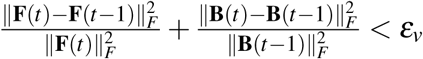 and 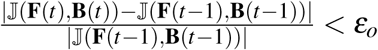, where *ε v* > 0, *ε*_*o*_ > 0 are pre-defined small constants. Since the objective function in (3) is non-convex and non-smooth, the SSEMQ algorithm can not guarantee to converge to the global minimal. However, we prove in the supplementary text that SSEMQ always converges to a stationary point of the objective function. The hyper parameters *ρ* and *λ* can be determined by Bayesian information criterion (BIC) with grid search, and expressions for their upper bounds are derived in the supplementary text. As mentioned earlier, if eQTLs are known and only eQTLs are included in the SEM (1) or equivalently (2), then term *ρ*‖**F**‖_1,**r**_ in (3) can be dropped, and the optimization problem (3) becomes the problem of GRN inference. We can easily modify (10) so that the SSEMQ algorithm is applicable to GRN inference. Note that SSEMQ employs a block coordinate descent (BCD) approach, because a column of **B** is estimated in (9), and a row of **F** is estimated in (11). It turns our that this BCD approach in conjunction with the proximal operator in (9) and (11) is much faster than the element-wise coordinate descent approach used by the SML algorithm (Cai *et al*., 2013).

## Computer Experiment

In this section, we conduct computer simulation studies to compare the performance of our SSEMQ algorithm with that of eight existing methods. These methods include MatrixEQTL-Linear (Shabalin, 2012), MatrixEQTL-ANOVA (Shabalin, 2012), GMAC (Yang *et al*., 2017), ElasticNet (Friedman *et al*., 2010), SIOL (Lee and Xing, 2012), GFLasso (Kim *et al*., 2009), LORs (Yang *et al*., 2013) and MTLasso2G (Chen *et al*., 2012). MatrixEQTL-Linear and MatrixEQTL-ANOVA test the association between each SNP and each gene with a linear regression model and an ANOVA model, respectively; they can find both *cis*-eQTLs and *trans*-eQTLs. GMAC applies a mediation test to the results of MatrixEQTL to determine *trans*-eQTLs mediated by *cis*-eGenes. Elastic Net, SIOL, GFLasso, LORs, and MTLasso2G all employ a linear regression model **Y** = **FX** + *µ***1**^*T*^ + **E**, where **Y** = [**y**_1_, **y**_2_, …, **y**_*N*_] is an *K* × *N* matrix of gene expression level of *N* samples and *K* genes, **X** is an *J* × *N* corresponding to the genotypes of *J* SNPs of *N* samples, and *µ* is a constant vector. eQTL mapping is performed by estimating **F**, through minimization of the following objective function: 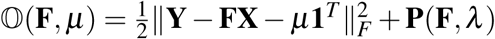, where the regularization term **P**(**F**, *λ*) is a function of **F** and hyperparameter *λ* and is specific to different methods. Apparently, ElasticNet, SIOL, GFLasso, LORS and MTLasso2G can find both *cis*- and *trans*-eQTLs. These methods use cross-validation to determine the optimal value of hyperparameter *λ*, and the SNPs corresponding to the nonzero entries of **F** are then identified as eQTLs. MatrixEQTL-Linear and MatrixEQTL-ANOVA output an p-value for the association between each SNP and each gene, and eQTLs are determined with a cutoff p-value. Here, we use a cutoff p-value such that the false discovery rate (FDR) is controlled at 10^−4^. Our SSEMQ employs BIC to determine hyperparameters *λ* and *ρ*, and model parameters **B** and **F** are estimated at the optimal values of *λ* and *ρ*.

### Algorithm 1 Sparse SEM based eQTL mapping (SSEMQ)

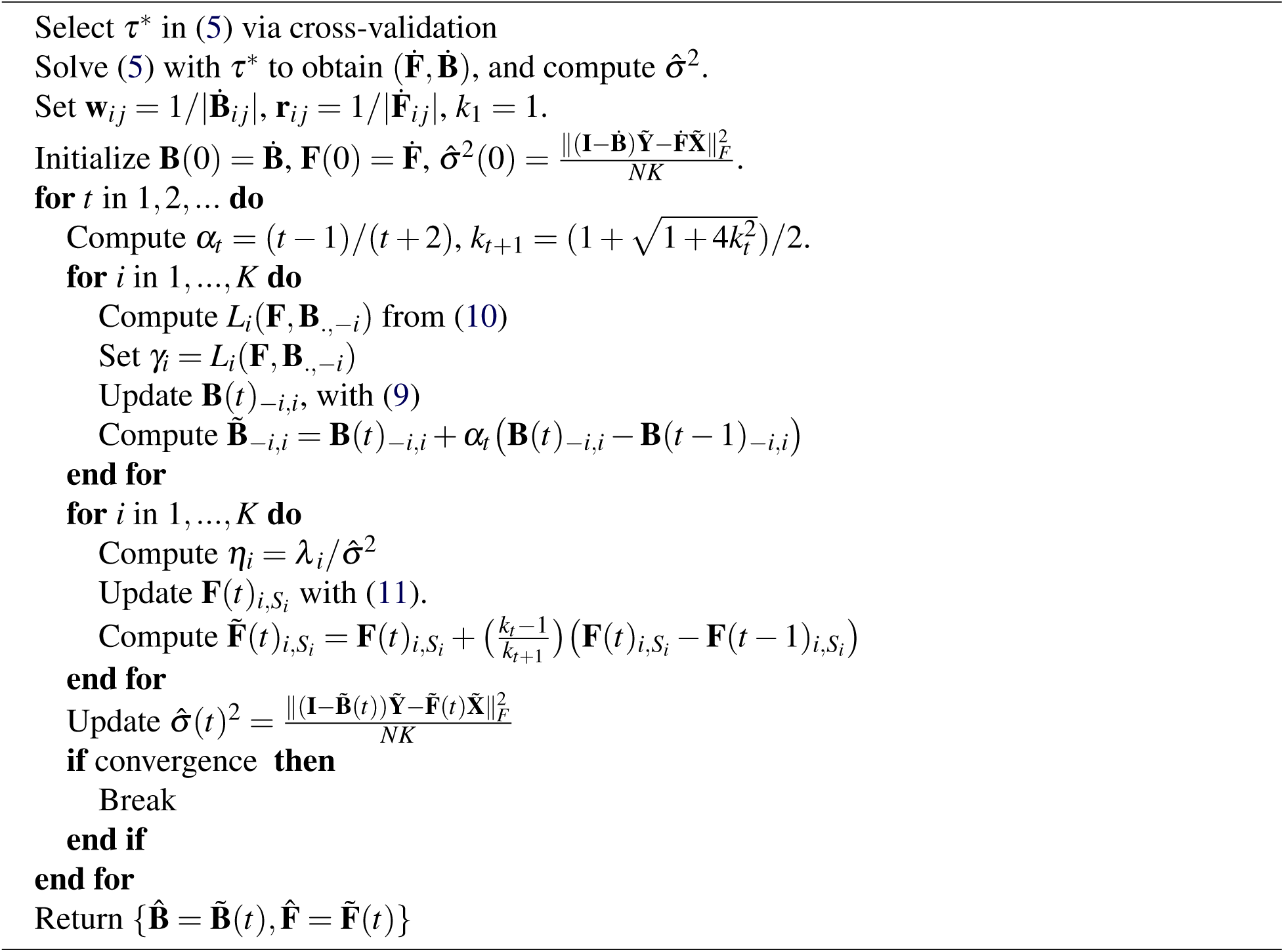

### 3.1 Synthetic genotype and gene expression data

We first simulated a large chromosome covered by evenly distributed SNPs. The distance between the neighboring SNPs is 20 cM. Three SNPs were randomly selected from the 15 SNPs within 300 cM of a gene as *cis*-eQTLs of the gene. The genotype of all SNPs were generated from an F2 cross, that was simulated based on the Stahl model using the *R/qtl* software package (Broman *et al*., 2003). Values 0 and 2 were assigned to two homozygous genotypes, respectively and 1 to the heterozygous genotype; and the genotype data matrix **X** of SNPs was generated. The effect sizes of eQTLs or the nonzero entries of matrix **F** were generated from a random variable uniformly distributed over interval [0.5, 5] or [−5, −0.5]. The minor allele frequencies (MAF) of these SNPs were chosen to be ≥ 0.10. We simulated both directed acyclic graph (DAG) and directed cyclic graph (DCG) for GRNs. Specially, the adjacency matrix **A** of a DAG or DCG of 30 or 100 genes with an expected number of edges per gene *n*_*d*_ = 1 was generated for the GRN, and the corresponding network coefficient matrix **B** was generated from **A** as follows. For **A**_*i j*_ = 0, we set **B**_*i j*_ = 0; and for **A**_*i j*_ = 1, we generated **B**_*i j*_ from a random variable uniformly distributed over interval [0.2, 0.6] or [−0.6, −0.2]. Error term **E** was independently sampled from the Gaussian distribution with zero mean and variance *σ* ^2^**I**, where *σ* ^2^ *∈* {0.10, 0.25}; *µ* was set to zero, and the sample size *N* varied from 80 to 500. Finally, simulated gene expression data matrix **Y** was obtained as **Y** = (**I** − **B**)^−1^(**FX** + **E**).

### 3.2 Accuracy of predicted *cis*-eQTLs

For each network configuration, 20 replicates of **X, F** and **B** were simulated, which were used to generate **Y**. Our SSEMQ algorithm and other seven methods mentioned earlier, excluding GMAC, were applied to **Y** and **X** to determine eQTLs. SSEMQ includes all SNPs within 300 cM of genes and outputs a set of *cis*-eQTLs based on the nonzero entries of the estimated **F**. For other seven methods, predicted eQTLs that are within 300 cM of a gene were identified as *cis*-eQTLs. The power of detection (PD) and the false discovery rate (FDR) were obtained by averaging the result from 20 replicates of the simulated data.

Figure 2 depicts the PD and FDR of predicted *cis*-eQTLs for the GRN of *K* = 100 genes. It is observed that the PD of our SSEMQ is much better than that of MatrixEQTL-Linear, MatrixEQTL-ANOVA, SIOL and MTLasso2G, and it is similar to that of Elastic Net, GFLasso and LORS. The FDR of SSEMQ is much smaller than that of all other seven methods across all sample sizes tested. The PD and FDR of *cis*-eQTLs for other DAG and DCG network configurations are shown in Figures S1-S3 and Figures S4-S6, respectively. Observations similar to those in Figure 2 can be seen from those figures. Interestingly, our SSEMQ and all the other methods based on penalized multiple linear regression outperform the single SNP methods, MatrixEQTL-Linear and MatrixEQTL-ANOVA, in terms of both PD and FDR. A similar observation was seen in (Lee and Xing, 2012).

**Figure 2.**
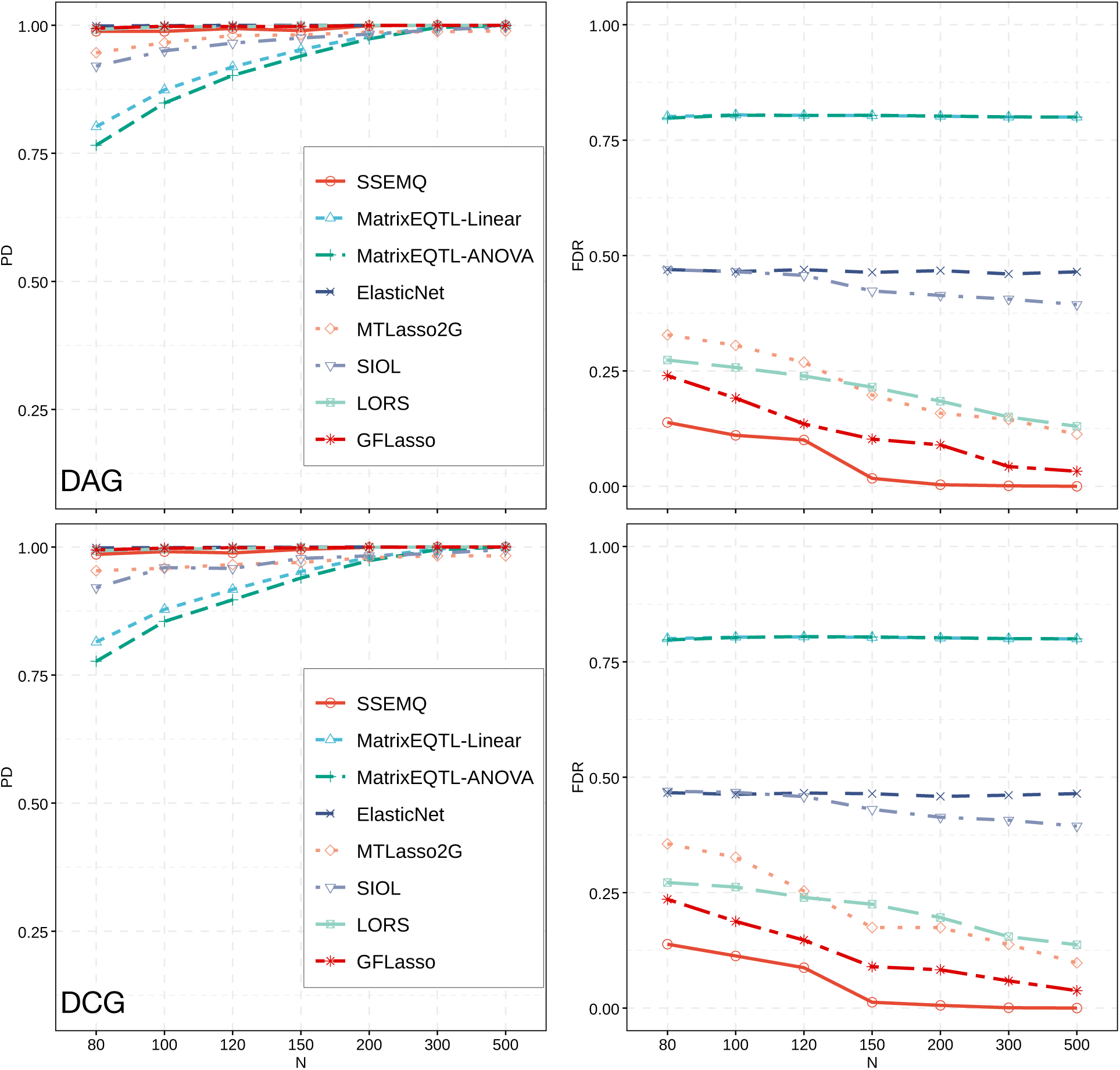
PD and FDR of *cis*-eQTLs for DAG (top) and DCG (bottom) with *K* = 100 genes. The noise variance *σ* ^2^ = 0.25. PD and FDR were obtained from 20 network replicates.

We further compare the performance of different methods using precision and recall (PR) curves. Possible *cis*-eQTLs were ranked according to their p-values for MatrixEQTL-Linear and MatrixEQTL-ANOVA, the absolute values of nonzero entries of **F** in SSEMQ, and the absolute values of regression coefficients for other methods. Precision and recall of *cis*-eQTLs were then calculated from the ranked SNPs under different cutoffs for SNPs to be considered as predicted *cis*-eQTLs. PR curves of predicted *cis*-eQTLs are shown in Figure 3 for the case *σ* ^2^ = 0.25 and in Figure S7 for the case *σ* ^2^ = 0.1. The area under PR Curve (AUPRC) for each method is listed in Table 1. It is observed that SSEMQ outperforms all other methods for all simulation settings, and that the single SNP methods, MatrixEQTL-Linear and MatrixEQTL-ANOVA, offer much worse performance than other multiple SNP methods.

**Table 1.** AUPRC of predicted eQTLs. The number of samples *N* = 80 and noise variance *σ* ^2^ = 0.10 and 0.25. The GRN has *K* = 100 genes with a DAG or DCG structure.

| Algorithm | $\sigma^2$ | *cis*-eQTLs | | *trans*-eQTLs | |
| --- | --- | --- | --- | --- | --- |
|  |  | DAG | DCG | DAG | DCG |
| SSEMQ | 0.10 | **0.998** | **0.998** | **0.887** | **0.856** |
|  | 0.25 | **0.990** | **0.988** | **0.867** | **0.807** |
| GMAC | 0.10 | – | – | 0.353 | 0.353 |
|  | 0.25 | – | – | 0.340 | 0.337 |
| MatrixEQTL<sub>Linear</sub> | 0.10 | 0.384 | 0.380 | 0.291 | 0.285 |
|  | 0.25 | 0.382 | 0.382 | 0.293 | 0.282 |
| MatrixEQTL<sub>ANOVA</sub> | 0.10 | 0.384 | 0.379 | 0.291 | 0.285 |
|  | 0.25 | 0.382 | 0.382 | 0.293 | 0.282 |
| ElasticNet | 0.10 | 0.967 | 0.969 | 0.330 | 0.283 |
|  | 0.25 | 0.963 | 0.963 | 0.285 | 0.242 |
| MTLasso2G | 0.10 | 0.900 | 0.925 | 0.201 | 0.206 |
|  | 0.25 | 0.920 | 0.865 | 0.215 | 0.219 |
| SIOL | 0.10 | 0.875 | 0.881 | 0.320 | 0.273 |
|  | 0.25 | 0.873 | 0.878 | 0.318 | 0.274 |
| LORS | 0.10 | 0.965 | 0.967 | 0.313 | 0.272 |
|  | 0.25 | 0.960 | 0.961 | 0.280 | 0.246 |
| GFLasso | 0.10 | 0.960 | 0.963 | 0.314 | 0.273 |
|  | 0.25 | 0.962 | 0.962 | 0.280 | 0.254 |

**Figure 3.**
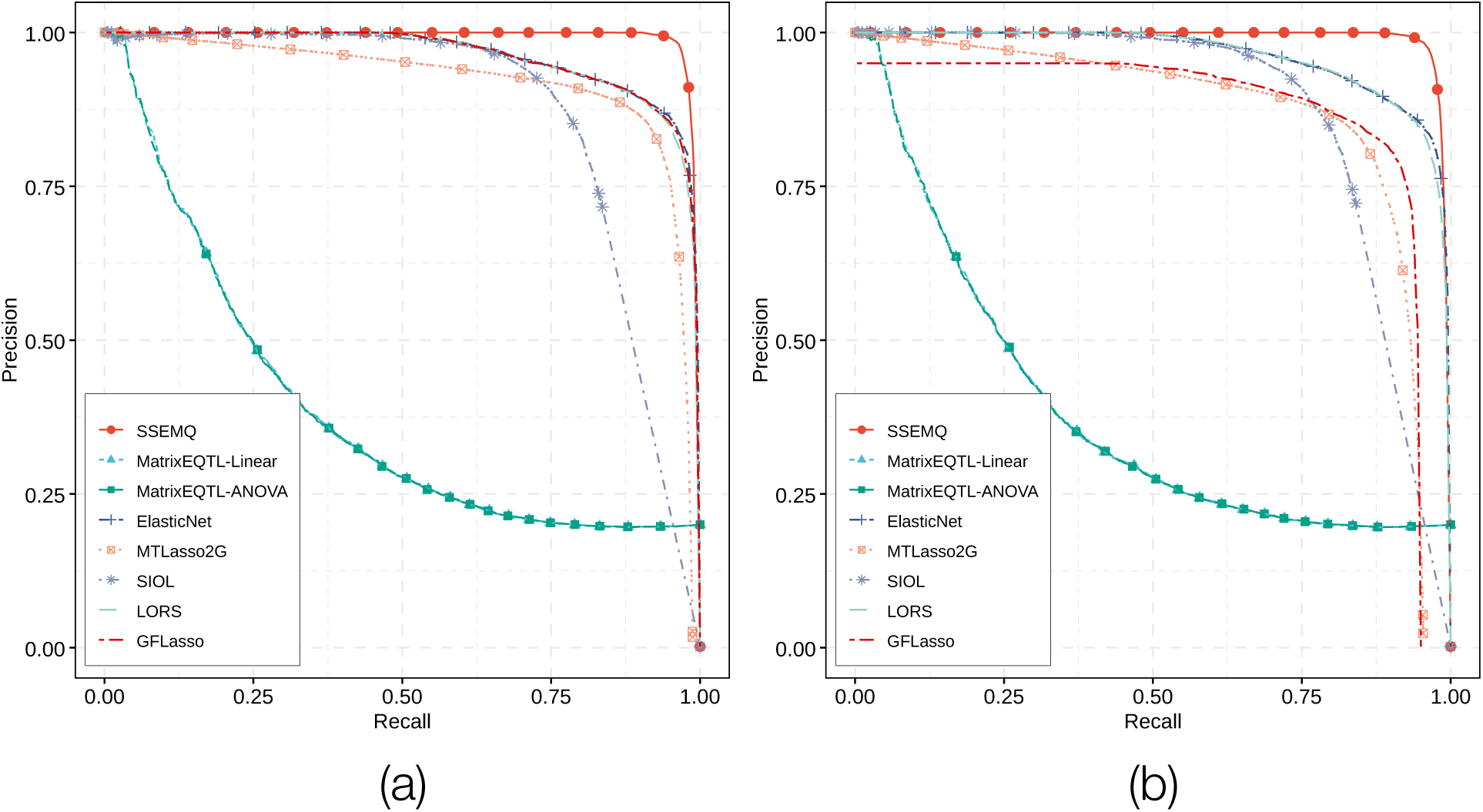
Precision-Recall curves of predicted *cis*-eQTLs. The number of samples *N* = 80 and noise variance *σ* ^2^ = 0.25. The GRN has *K* = 100 genes with a DAG (a) or DCG (b) structure.

### 3.3 Accuracy of predicted *trans*-eQTLs

While matrix **F** estimated by our SSEMQ algorithm yields *cis*-eQTLs, we can also obtain *trans*-eQTLs from **F** and **B** as follows. From SEM in (1), we have **y**_*i*_ = (**I** −**B**)^−1^**Fx**_*i*_ + (**I** −**B**)^−1^(*µ* + *ε*_*i*_), *i* = 1, …, *N*. If we define matrix **G** = (**I** −**B**)^−1^**F**, it is clear nonzero entries of the *k*th row of **G** gives all eQTLs of gene *k*, and the *trans*-eQTLs of gene *k* are then obtained by excluding the *cis*-eQTLs that have been identified from **F**. GMAC determines the *trans*-eQTLs with a mediation test based on *cis*-eQTLs, *cis*-eGenes and *trans*-eGenes identified with MatrixEQTL (Shabalin, 2012). For other seven methods, predicted eQTLs that are not located within 300 cM of a gene were designed as *trans*-eQTLs.

Figure 4 and S8 show the PR curves of predicted *trans*-eQTLs for *σ* ^2^ = 0.25 and 0.1, respectively. Table 1 lists the AUPRCs calculated from PR curves in Figure 4 and S8. It is seen from these results that our SSEMQ algorithm outperforms significantly all other methods. The performance of GMAC is slightly better than MatrixEQTL as expected. The performance of SSEMQ and MatrixEQTL for *trans*-eQTLs is slightly worse than that for *cis*-eQTLs, whereas the performance of other five methods for *trans*-eQTLs is significantly worse than that for *cis*-eQTLs. Performance of all methods is slightly better when the GRN is DAG than when the GRN is a DCG.

**Figure 4.**
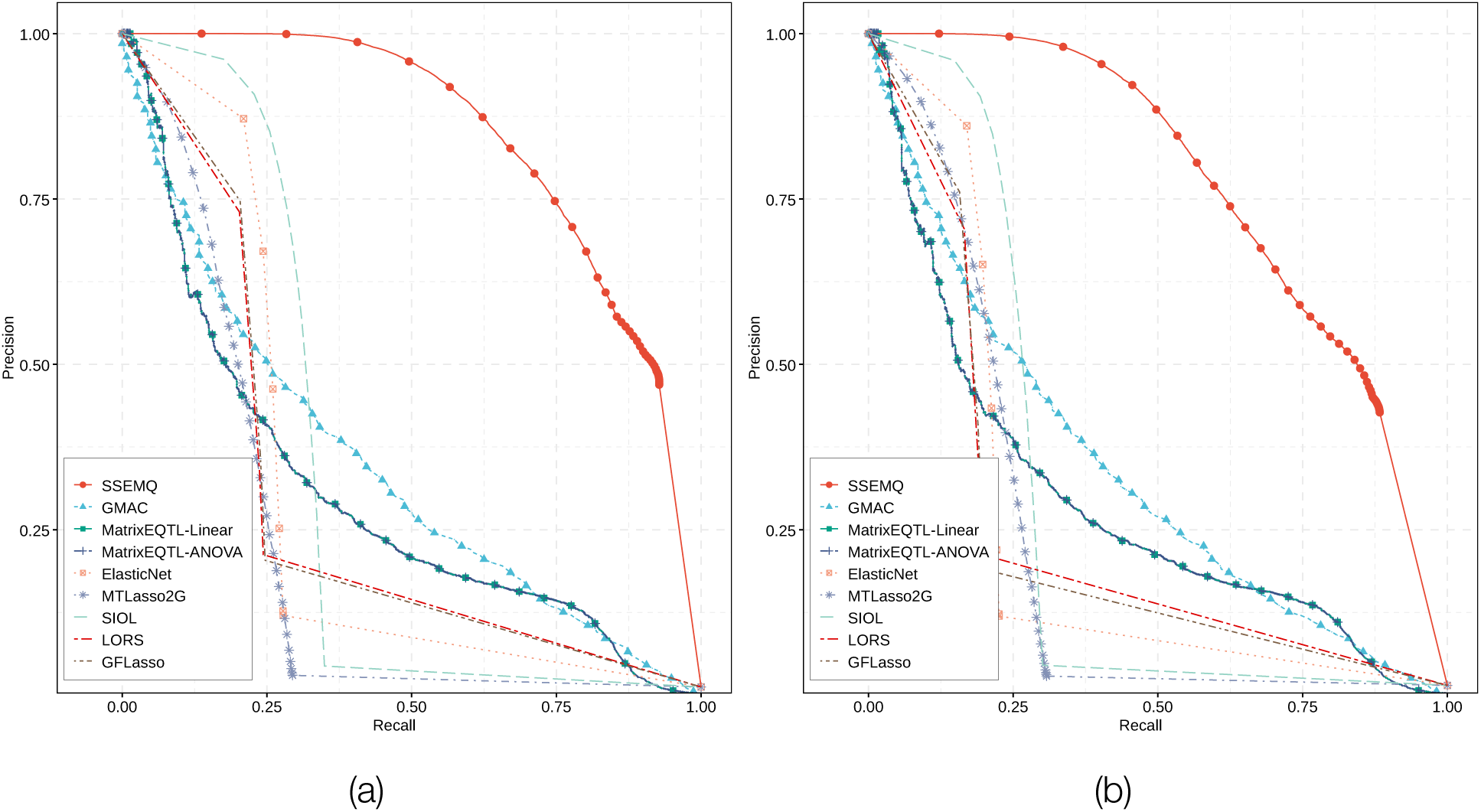
Precision-Recall curves of predicted *trans*-eQTLs. The number of samples *N* = 80 and noise variance *σ* ^2^ = 0.25. The GRN has *K* = 100 genes with a DAG (a) or DCG (b) structure.

Of note, for our SSEMQ algorithm, there are *P*_*c*_ = 1, 500 unknown entries in **F** and *P*_*t*_ = *K*^2^ − *K* = 9, 900 unknown entries in **B** to be estimated, when the GRN contains *K* = 100 genes. For other methods based on multiple linear regression, the **F** matrix has *P* = 100 × 1500 = 15, 000 unknown. Therefore, the number of unknown coefficients to be estimated in SNP analysis is much greater than the number of data sample *N*. Due to the regularization technique used in SSEMQ and multiple regression methods, our SSEMQ algorithm and these multiple SNP methods can handle the problem of “large *p* and small *N*”.

### 3.4 Accuracy of GRN inference

Assuming that eQTLs are known, we employed SEM to infer one GRN (Cai *et al*., 2013) or jointly infer two GRN (Zhou and Cai, 2020). Here eQTLs are unknown and our main purpose is eQTL mapping, although the GRN is also inferred as a byproduct. In fact, GRN inference helps to identify trans-eQTLs. We have shown the performance of SSEMQ on eQTL-mapping in previous two sections. Here we will show the performance of SSEMQ in GRN inference. As described earlier, we simulated genotypes **X** of a set of SNPs, and the adjacency matrix **A** of the GRN, and then generated network matrix **B** and gene expression data **Y**. We applied SSEMQ to the simulated data **X** and **Y**, and estimated 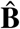. Setting the nonzero entries of estimated 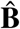 to one, we obtained estimated adjacency matrix **Â** of the GRN. Comparing **A** and **Â**, we obtained PD and FDR of detected edges of the GRN, which are shown in Figure S9 for DAG networks and in Figure S10 for DCG networks. Both figures show that the PD is close to one and FDR is very small, close to zero especially when sample size *N ≥* 150.

## Real data analysis

We applied our SSEMQ algorithm to perform eQTL mapping with the genotype and gene expression data of human breast tissues from the GTEx Project v6p release (Lonsdale *et al*., 2013). This dataset contains expression levels of 24, 283 genes and genotypes of 11, 959, 411 unique SNPs in 183 individuals. To obtain informative genes for QTL mapping, we calculated the coefficient of variation (CoV) of the RPKM expression values of each gene in 183 samples. Genes with an average RPKM < 0.1 and CoV < 1.0 were excluded, which resulted in a set of 884 protein coding genes. Then, the **Y** matrix of the SEM (1) was formed with log-transformed and quantile-normalized RPKMs (Lonsdale *et al*., 2013) of the 884 genes across 183 samples. SNPs with a minor allele frequency (MAF) of 5% or less were removed. SNPs located within 1M base pairs (bps) to each of the 884 protein coding genes were extracted. GTEx has performed *cis*-eQTL mapping with FastQTL (Ongen *et al*., 2016) and provided p-values for all SNPs. Retaining those SNPs with a p-value < 0.10, we obtained 23, 934 SNPs as candidate *cis*-eQTLs for the 884 genes. The genotypes of these SNPs were transformed to values {0, 1, 2} using the following mappings: 0*/*0 → 0, 0*/*1 → 1 and 1*/*1 → 2; these values were used to form the **X** matrix in the SEM (1).

We then applied the SSEMQ algorithm to the data matrices **X** and **Y** to obtain estimates of **B** and **F**, 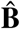 and 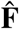. To find *cis*- and *trans*-eQTLs more reliably, we employed the stability selection technique (Meinshausen and Bühlmann, 2010) to determine the nonzero entries of 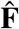 and 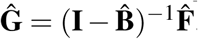. Stability selection was described in the Supplementary text. Pairs of *cis*-eQTL and *cis*-eGenes were identified from the nonzero entries of 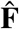, while pairs of *trans*-eQTL and *trans*-eGenes were determined from the nonzero entries of **Ĝ**. In total, we found 3, 239 eQTL/eGene pairs, among which 2, 048 are *cis*-eQTL/*cis*-eGene pairs of 752 genes, and 1, 191 are *trans*-eQTL/*trans*-eGene pairs of 464 genes. We found that 92.58% (699/755) of the significant *cis*-eQTL-*cis*/eGene pairs detected by GTEx at FDR < 0.1 (p-value ¡0.0007) were included in our *cis*-eQTL-*cis*/eGene pairs.

An eQTL that is associated with a relatively large number of genes is referred to as a hot-spot (Yang *et al*., 2013). Since all *trans*-eQTLs and *trans*-eGenes identified here are mediated by *cis*-eGenes, we refer *cis*-eGene and its *cis*-eQTLs that are associated with a large number of *trans*-eGenes to as a hot-spot. We performed gene ontology (GO) enrichment analysis for the set of genes associated with each hot-spot. The top 30 hot-spots with largest number of genes and significantly enriched GO terms (adjusted p-value < 10^−5^) are listed in supplementary table S1. The number of *trans*-eGenes associated with these 30 hot-spots is ≥6. The first hot-spot contains four *cis*-eQTs (4 72208873 G A b37, 4 71469529 C T b37, and 4 71219731 G T b37, 4 71226021 A G b37), *cis*-eGene JCHAIN, and 27 downstream *trans*-eGenes. We extracted the sub-network of JCHAIN and its 27 downstream genes determined by 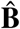, and plotted the network in Figure 5. The top ten genes with highest degrees in the network are SYT8, OXTR, SLC34A2, FOLR1, PIGR, MZB1, KLK5, MMP7, and ELF5. Interestingly, six of these ten genes were found to be related to immunity. JCHAIN gene encodes the joining chain of polymeric IgA and IgM (Max *et al*., 1986). SLC34A2 was reported to induce abnormal alveolar type II (AT II) cells to transform into cancer cells, and the AT II cells possess immune functions of synthesizing factors of immune regulation (Yang *et al*., 2014). FOLR1 is known as a tumor-associated antigen (Lu *et al*., 2004; Lu and Low, 2003), which helps immunoglobulins target tumor cells and stimulates immune response. PIGR is a polymeric immunoglobulin receptor gene; it binds polymeric IgA and IgM and transports them from the basolateral surface of the epithelium to the apical side (Turula and Wobus, 2018). MZB1 is expressed in innatelike B cells, and is involved in the regulation of IgM and IgA’s response (Suzuki *et al*., 2019). ELF5 gene is a key transcriptional determinant of tumor subtypes and has been found to help recognition of antibodies (Piggin *et al*., 2020).

**Figure 5.**
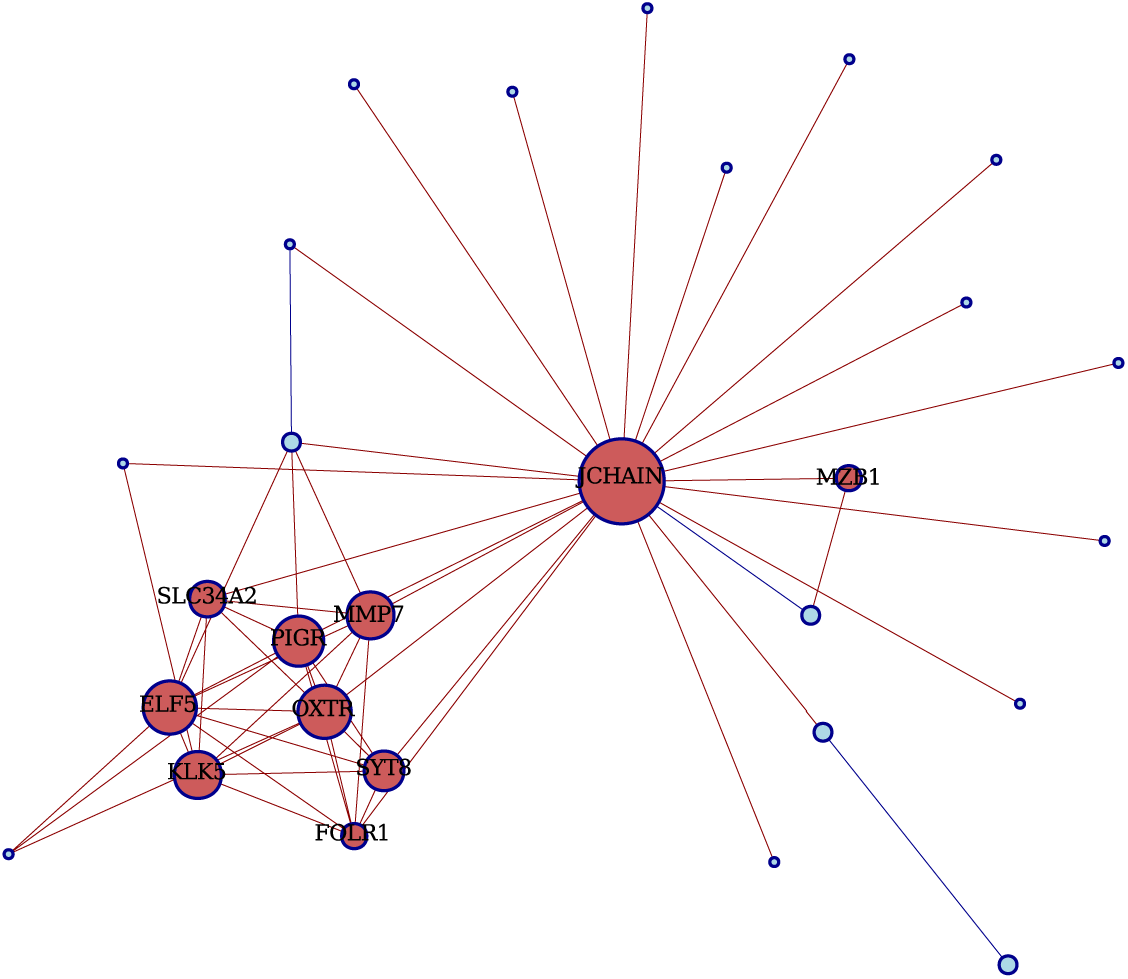
The sub-network of 28 genes associated with the first hot-spot that includes the mediator eGene JCHAIN and four *cis*-eQTLs (4 72208873 G A b37, 4 71469529 C T b37, 4 71219731 G T b37, 4 71226021 A G b37). The size of a gene node is proportional to its degree, the number of edges that the gene connects to. The top 10 genes with highest degrees are labeled.

## Conclusion

In this paper, we have developed an efficient algorithm, named SSEMQ, for eQTL mapping. The algorithm outputs *cis*-eQTL and *cis*-eGene pairs and the GRN structure, which are further used to determine *trans*-eQTL and *trans*-eGene pairs mediated by *cis*-eGene pairs. Computer simulations showed that SSEMQ offered better performance on identifying both *cis*- and *trans*-eQTLs than eight existing methods. The superior performance on *cis*-eQTL mapping is due to the fact that correlated expression levels of genes influenced by the same eQTL is exploited jointly, while the superior performance on *trans*-eQTL mapping is due to the GRN inference. The SSEMQ algorithm was further employed to analyze a real dataset of human breast tissues, yielding a number of eQTLs and eGenes. GO enrichment analysis of the networks of eGenes shows that genes influenced by the same set of eQTLs may be involved in certain common biological processes and functions.

## Supplementary Text S

### Hyperparameter selection

We use *K*-fold cross-validation (CV) to determine the value of *τ* for ridge regression (5) and values of *λ* and *ρ* for SSEMQ, where *K* typically equals to 5 or 10. We search *τ* over a sequence of 50 values increasing from 10^−6^ to 10^2^ evenly on the logarithm scale, and the optimal value of *τ* is chosen to minimize the predication error calculated from the test data. We employ a grid search strategy to determine the optimal values of *λ* and *ρ*. We first determine the maximum value of *λ*, namely *λ*_max_, then choose a set of *k*_1_ values for *λ*, denoted as sequence 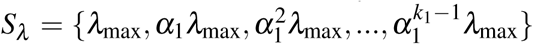, where 0 < *α*_1_ < 1. For each value of *λ ∈ S*_*λ*_, we find the maximum value of *ρ*, namely *ρ*_max_(*λ*), and then choose a set of *k*_2_ values for *ρ*, denoted as 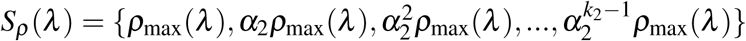, where 0 < *α*_2_ < 1. This gives a set of *K* = *k*_1_*k*_2_ pairs of (*λ, ρ*), and Bayesian information criterion (BIC) is used to tune hyperparameters (*λ, ρ*). The optimal values of *λ* and *ρ* are chosen to minimize the BIC scores.

Next, we derive the maximum values of *λ* and *ρ*. The value *λ*_max_ yields **B** = **0**, and can be found from the result in (Cai *et al*., 2013) as follows:

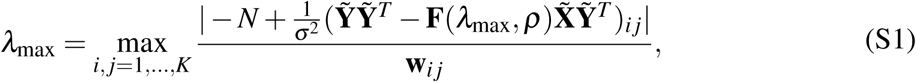

where **F**(*λ*_max_, *ρ*) is obtained through (11) iteratively after setting 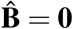. The maximum value of *ρ* at a given *λ* value, denoted as *ρ*_max_(*λ*), yields **F** = **0**; and *ρ*_max_(*λ*_max_) can be found as follows:

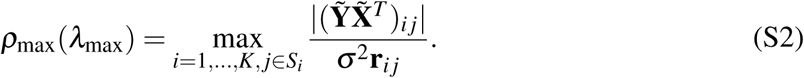

For each *λ* in *S*_*λ*_, we obtain *ρ*_max_(*λ*) as follows:

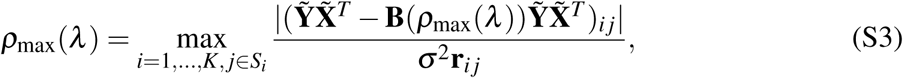

where **B**(*ρ*_max_(*λ*)) can be obtained from (4) with **F** = **0** or equivalently the following optimization problem:

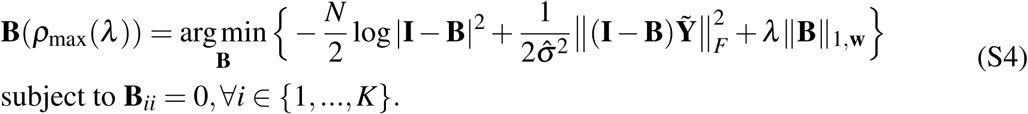

When the number of unknown is large, tuning hyperparameters with BIC is generally faster than the CV. Therefore, the hyperparameters (*λ, ρ*) of SSEMQ are selected by minimizing the following BIC score:

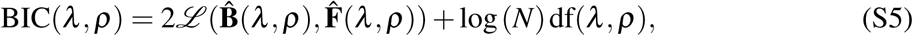

where 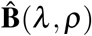 and 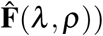 are estimates of **B** and **F** at 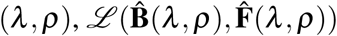 is the negative log-likelihood defined in (2), and df(*λ, ρ*) is the total number of nonzero elements in 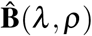 and 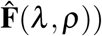.

### Convergence analysis

When the objective function in an optimization problem is non-convex and non-smooth, it is possible that the coordinate descent method fails to converge. We next prove that the SSEMQ algorithm converges to a stationary point, because the objective function satisfies the conditions for the convergence of the PALM method specified in (Bolte *et al*., 2014). Specifically, we prove 𝕁 (**F, B)** in (8) has the following properties:

1. inf ℍ(**F, B**) > −∞, when **B** *∈* dom ℍ = {**B** : det(**I**−**B**) ≠ 0}, inf *f*_*i*_(**F**_*i*,._) > −∞ and inf *g*_*i*_(**B**_.,*i*_) > −∞, *i* = 1, …*K*
2. ∇ℍ(**B**_−*i,i*_), *i* = 1, …, *n*, is Lipschitz continuous with a Lipschitz constant *L*_*i*_(**F, B**_.,−*i*_) for **B** *∈* dom ℍ:

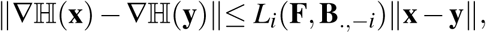

and ∇ℍ(**F**_−*i,S_i_*_), *i* = 1, …, *n* is Lipschitz continuous with a Lipschitz constant 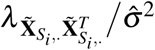.
3. ℍ(**F, B**) has continuous first and second derivatives when **B** *∈* dom ℍ.
4. 𝕁(**F, B**) satisfies the Kurdyka-Łojasiewicz(KL) property.

Note that properties 1-3 are identical to the properties in assumption B of (Bolte *et al*., 2014). Based on the result in (Stephen Boyd and Lieven Vandenberghe, 2004), these four properties guarantee that SSEMQ algorithm converges to a critical point. First, it is apparent that ℍ(**F, B**) > −∞ and therefore 𝕁(**F, B**) > −∞, *∀* **B** *∈* dom ℍ. Second, it is not difficult to show that ℍ(**F, B**) is differentiable w.r.t. **B**_*i*,−*i*_, *i* = 1, …, *n* and the first-order and second-order derivatives are continuous in dom ℍ. Therefore, property 3 is satisfied.

Third, we proved in next section that ∇ℍ(**B**_−*i,i*_) is Lipschitz continuous with constant *L*_*i*_(**F, B**_.,−*i*_) given in (10). Moreover, based on assumption B(iii) of (Bolte *et al*., 2014), *L*_*i*_(**F, B**_.,−*i*_) guarantees that proximal steps in the SSEMQ algorithm remain well-defined, because we have

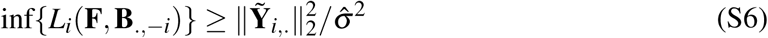

Finally, we proved that property 4 holds. The non-smooth functions *f*_*i*_(**F**_*i*,,_) and *g*_*i*_(**B**_.,*i*_) in 𝕁(**F, B**) are lasso penalty terms w.r.t. **F**_*i*,._ and **B**_.,*i*_, and they are semi-algebraic as shown in (Yangyang Xu and Wotao Yin, 2013). The *ℓ*_2_-norm 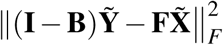 is apparently semi-algebraic. We next prove that 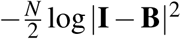 is semi-algebraic too. We can regard 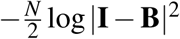 as a composite function of **B** as follows

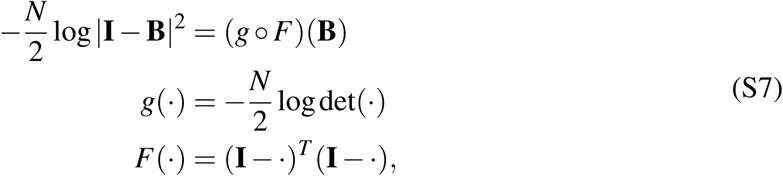

Function *g*(·) is locally convex function (Stephen Boyd and Lieven Vandenberghe, 2004). Based on the result in (Yangyang Xu and Wotao Yin, 2013), it is not difficult to show that *g*(·) satisfies the KL property, and it can be shown that function *F* : ℝ^*n*×*n*^ → ℝ^*n*×*n*^ is continuously differentiable in dom 𝕁. As all terms of 𝕁(**F, B**) are KL functions, the sum of theses KL functions should satisfy the KL property (Guoyin Li and Ting Kei Pong, 2017). This completes the proof that 𝕁(**F, B**) satisfies properties 1-4.

### Derivation of the Lipschitz constant of ∇ℍ(B−*i,i*)

Based on ℍ(**F, B**) defined in (8), it is not difficult to find 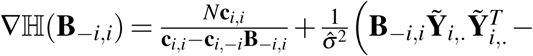 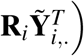, which has the following property:

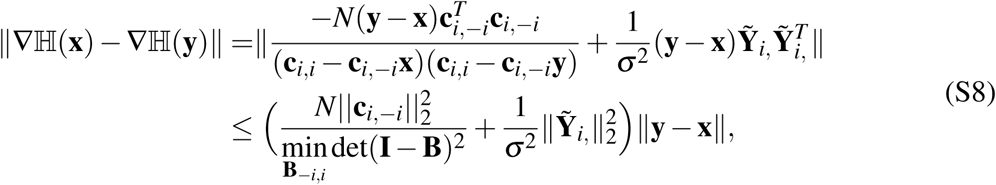

where 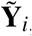, is the 1 × *N* vector for *i*-th roIw of 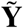. From (S8), it is apparent that Lipschitz constant 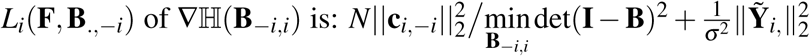.To calculate *L*_*i*_ (F, **B**_−.,*i*_), we need to minimize det(**I B**)^2^ w.r.t **B**_−*i,i*_, when **B**_., *j*_, *j* = 1, 2, …, *n, j* ≠ *i*, are fixed, which will be derived as follows.

Defining Θ = **I − B**, and let *θ* _.,*i*_ be the *i*-th column of Θ and Θ_., *i*_ be the sub-matrix of Θ that excludes *θ* _.,*i*_. Then, we can write det(Θ)^2^ as follows,

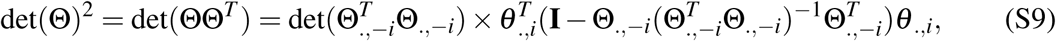

where Θ_−*i*,−*i*_ is the sub-matrix of Θ excluding the *i*th row and the *i*th column. Letting **b**_*i*_ = **B**_−*i,i*_, we can rewrite det() as

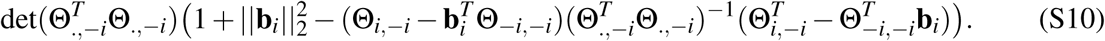

Minimizing det(Θ)^2^ in (S10) w.r.t. **b**_*i*_ yields

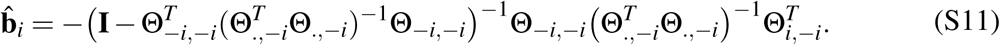

Substituting 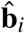 into (S10) gives the minimum value of det(Θ)^2^. In practice, to ensure numerical stability, we modify the 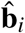 in (S11) as follows,

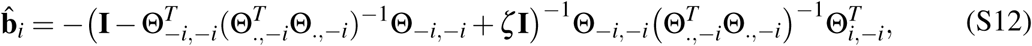

where *ζ* is a small positive constant. This modification can be regarded as minimizing det(Θ)^2^in (S12) subject to the constraint ‖ **b**_*i*_ ‖ _2_≤ *c*, where *c* is a positive constant. This is a reasonable assumption because in real GRNs, entries of **B** are bounded. In our implementation, we chose *ζ* = 10^−16^ and we did not observe any numerical instability in all of our numerical experiments.

### Stability selection

In real data analysis, we employed the stability selection technique (Meinshausen and Bühlmann, 2010) to determine reliably the nonzero entries of 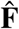 and 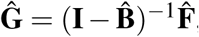, which were then used to determine *cis*- and *trans*-eQTLs. We first determined the optimal hyperparameter 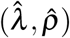 by minimizing the BIC score in (S5). We then randomly divided the dataset (**Y, X**) into two subset of equal and inferred 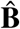 and 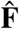 with our SSEMQ algorithm using each subset of the data and the optimal hyperparameter values 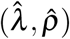. We repeated this process *N* times, which yielded 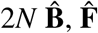, and **Ĝ**. We chose *N* = 50. Let *q*_*i j*_ and *t*_*i j*_ be the total number of nonzero 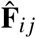 and **Ĝ** *i j*, respectively, in 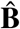 and 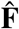 matrices. Then, *f* (**F**)_*i j*_ = *q*_*i j*_*/*(2*N*) and *f* (**G**)_*i j*_ = *t*_*i j*_*/*(2*N*) give the frequencies of nonzero 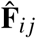 and **Ĝ** *i j*, respectively. Those 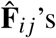 and **Ĝ** *i j*’s with *f* (**F**)_*i j*_ > *π*_thr_ and *f* (**G**)_*i j*_ > *π*_thr_, where *π*_thr_ is constant in the interval [0.6, 0.9] (Meinshausen and Bühlmann, 2010), were deemed to be nonzero. We used *π*_thr_ = 0.6 in our data analysis

## Supplementary Figure

**Figure S1.**
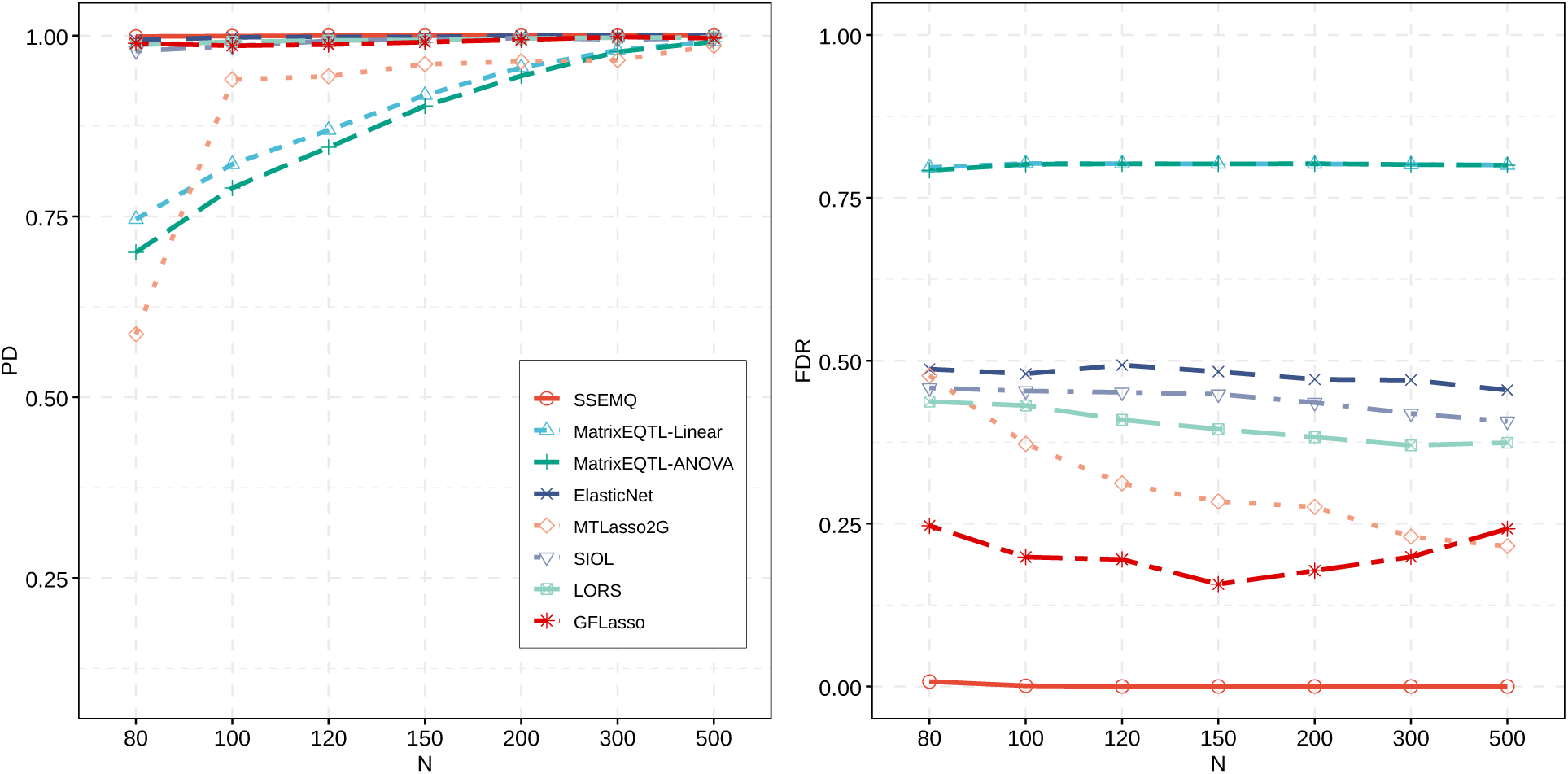
PD and FDR of *cis*-eQTLs for DAG with *K* = 30 genes. The number of samples *N* varies from 80 to 500 and noise variance *σ* ^2^ = 0.10. PD and FDR were obtained from 20 network replicates.

**Figure S2.**
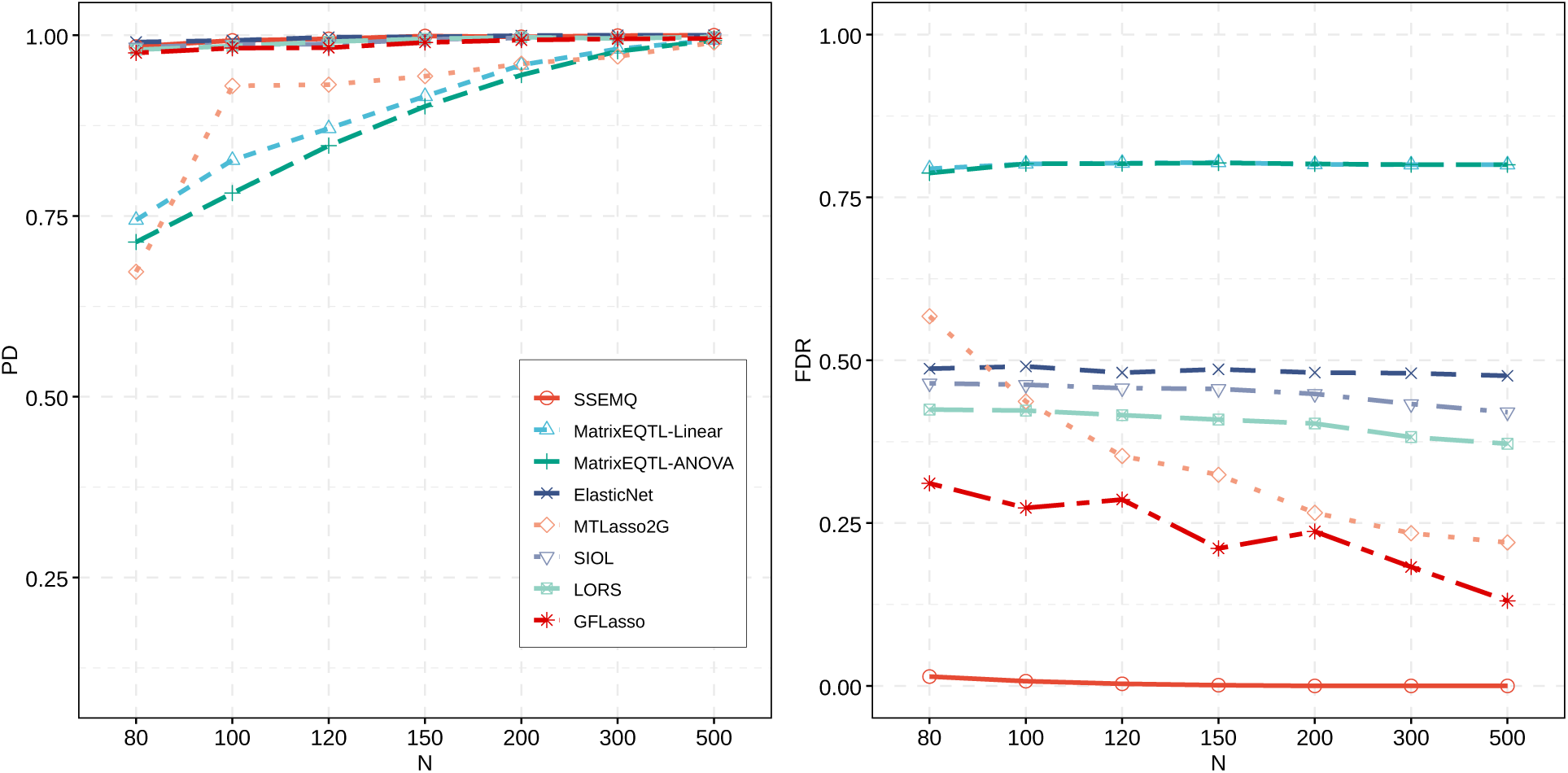
PD and FDR of *cis*-eQTLs for DAG with *K* = 30 genes. The number of samples *N* varies from 80 to 500 and noise variance *σ* ^2^ = 0.25. PD and FDR were obtained from 20 network replicates.

**Figure S3.**
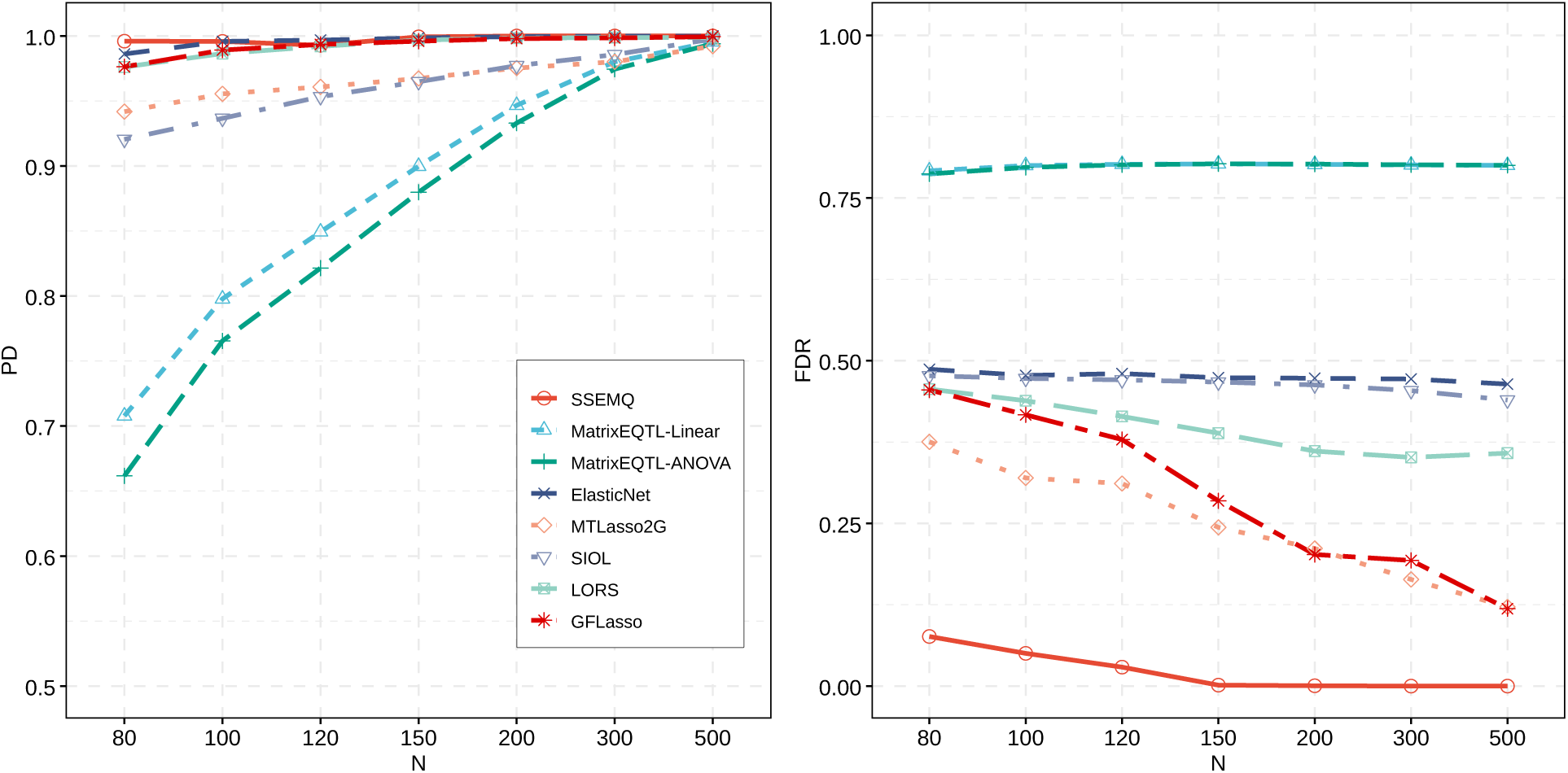
PD and FDR of *cis*-eQTLs for DAG with *K* = 100 genes. The number of samples *N* varies from 80 to 500 and noise variance *σ* ^2^ = 0.10. PD and FDR were obtained from 20 network replicates.

**Figure S4.**
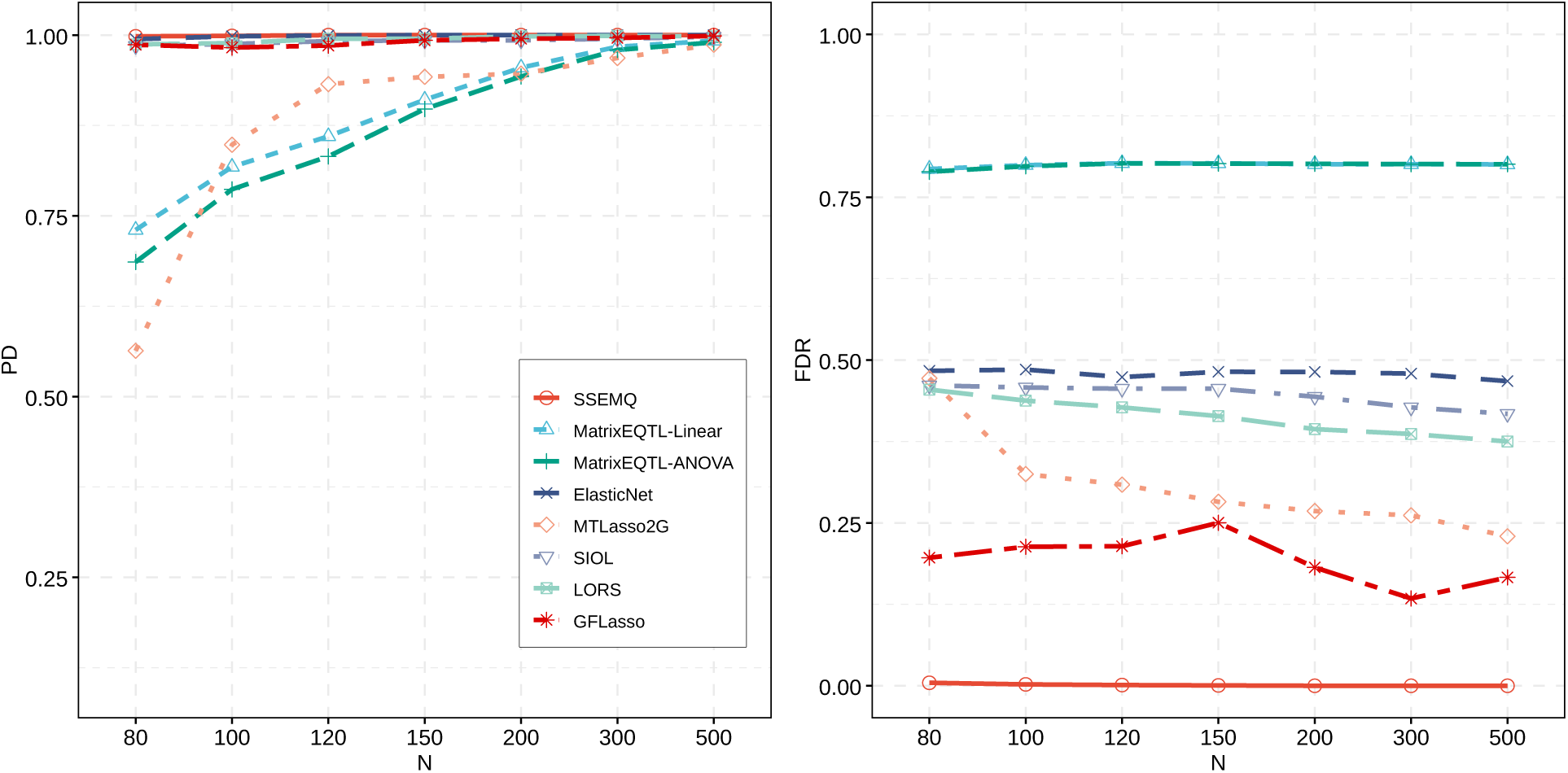
PD and FDR of *cis*-eQTLs for DCG with *K* = 30 genes. The number of samples *N* varies from 80 to 500 and noise variance *σ* ^2^ = 0.10. PD and FDR were obtained from 20 network replicates.

**Figure S5.**
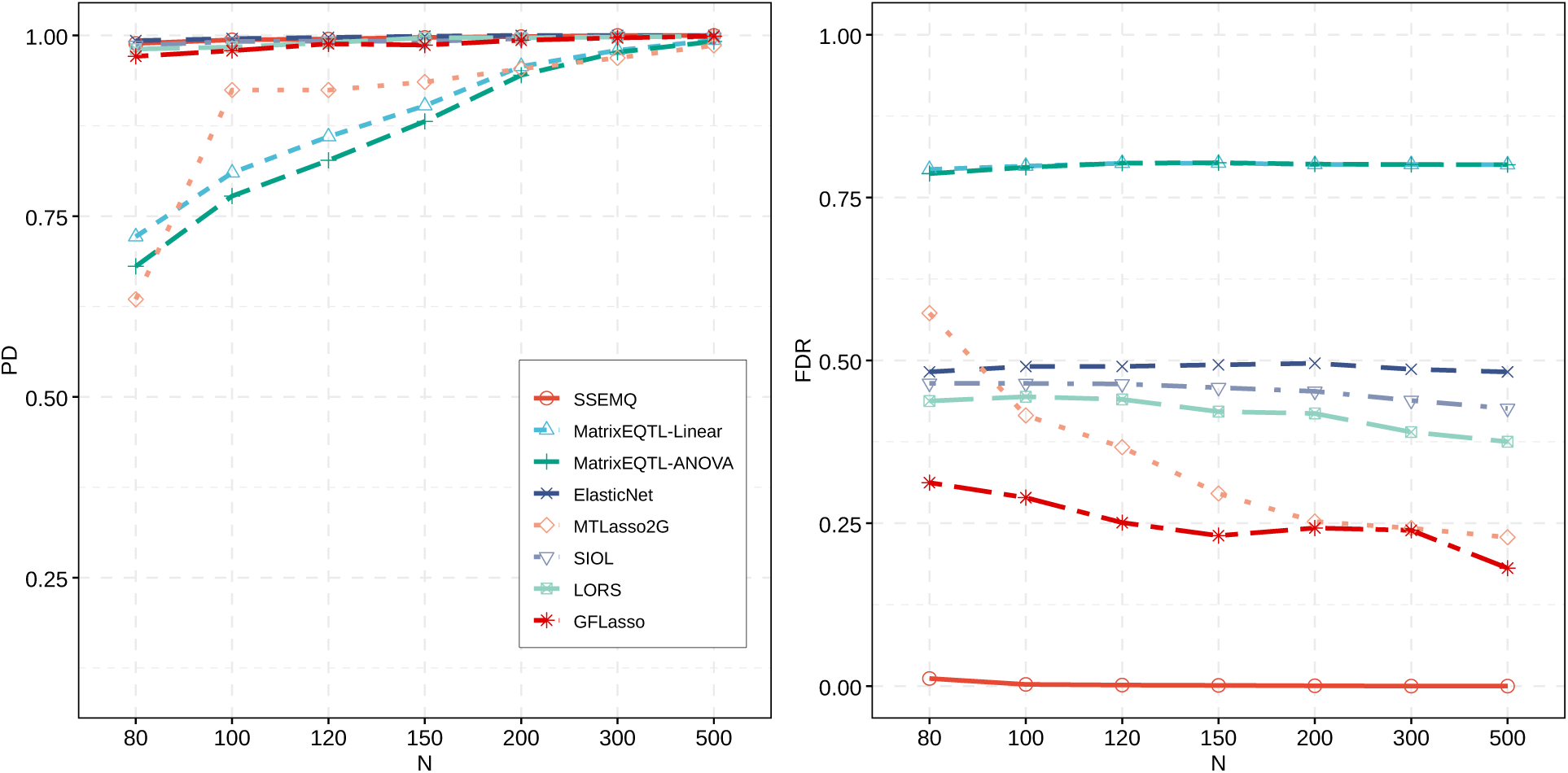
PD and FDR of *cis*-eQTLs for DCG with *K* = 30 genes. The number of samples *N* varies from 80 to 500 and noise variance *σ* ^2^ = 0.25. PD and FDR were obtained from 20 network replicates.

**Figure S6.**
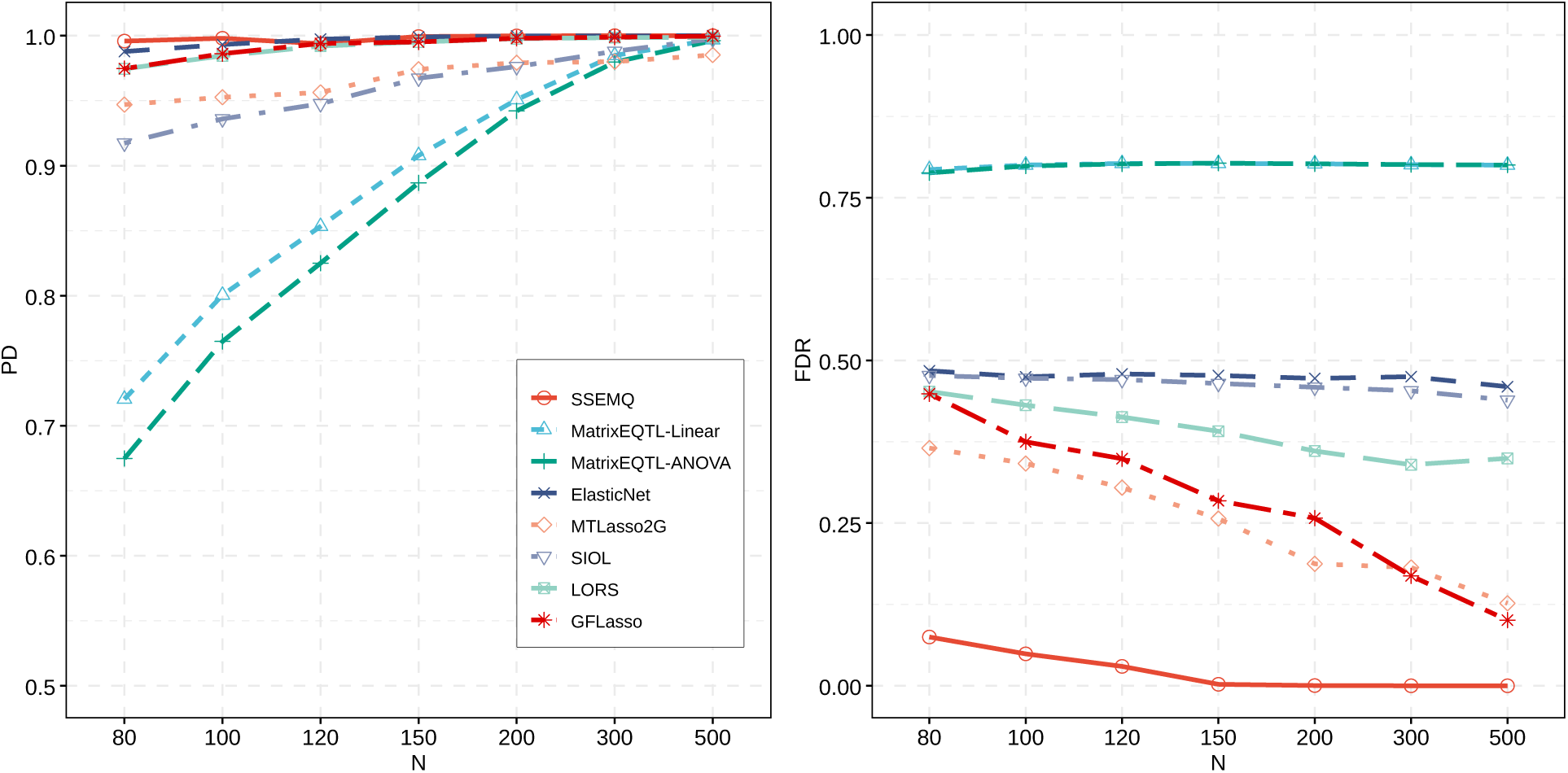
PD and FDR of *cis*-eQTLs for DCG with *K* = 100 genes. The number of samples *N* varies from 80 to 500 and noise variance *σ* ^2^ = 0.10. PD and FDR were obtained from 20 network replicates.

**Figure S7.**
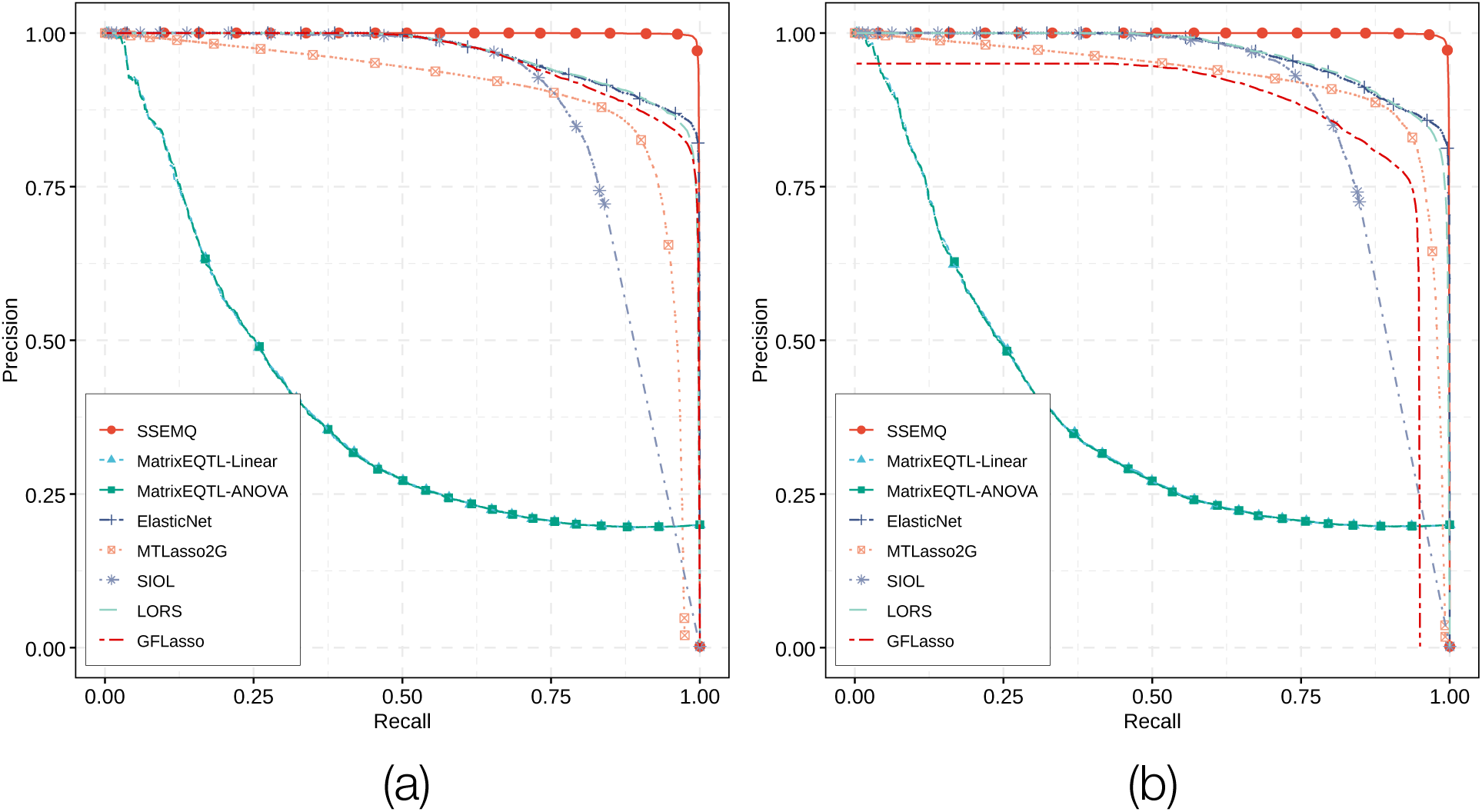
Precision-Recall curves of predicted *cis*-eQTLs. The number of samples *N* = 80 and noise variance *σ* ^2^ = 0.1. The GRN has *K* = 100 genes with a DAG (a) or DCG (b) structure.

**Figure S8.**
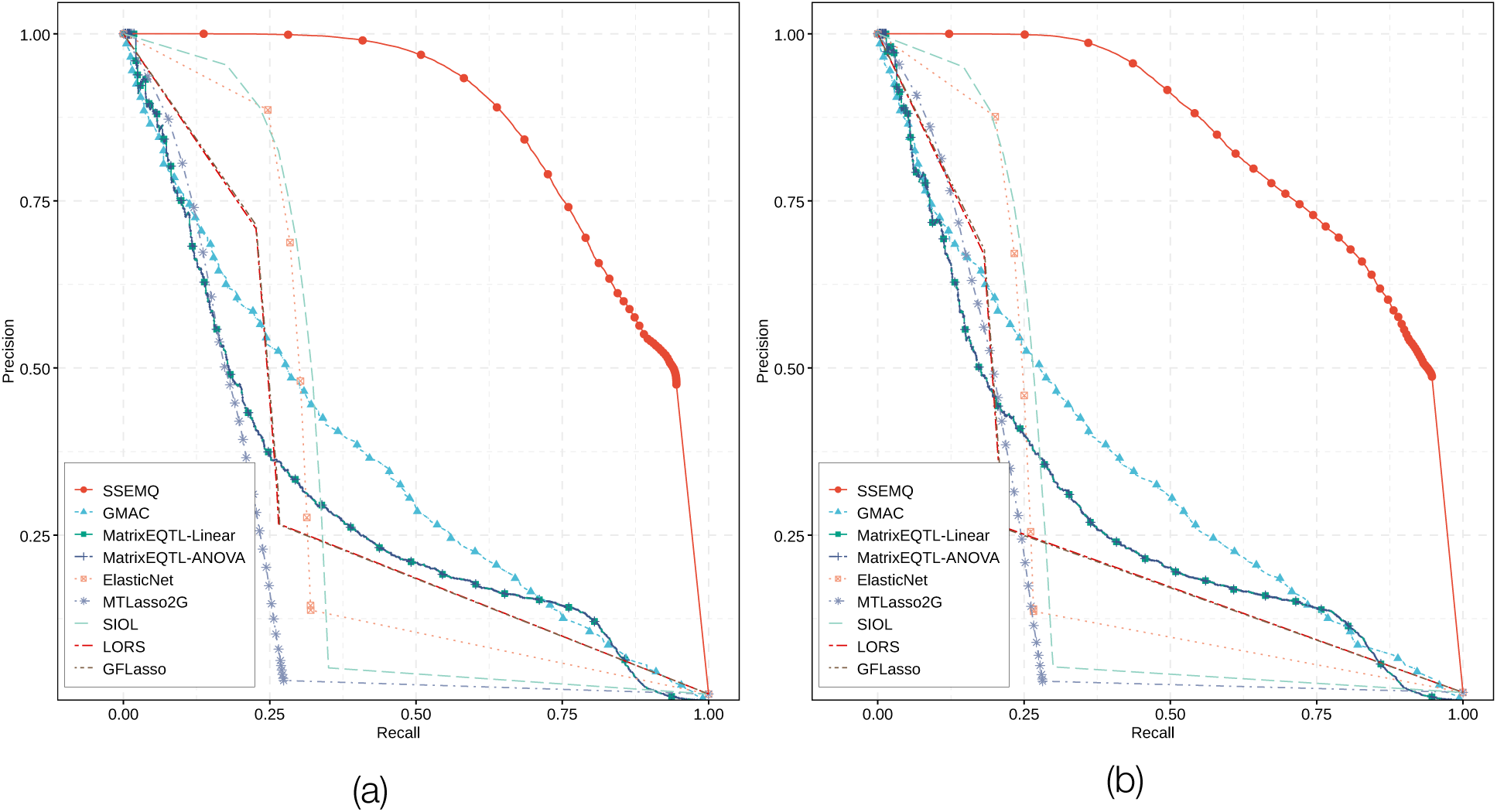
Precision-Recall curves of predicted *trans*-eQTLs. The number of samples *N* = 80 and noise variance *σ* ^2^ = 0.10. The GRN has *K* = 100 genes with a DAG (a) or DCG (b) structure.

**Figure S9.**
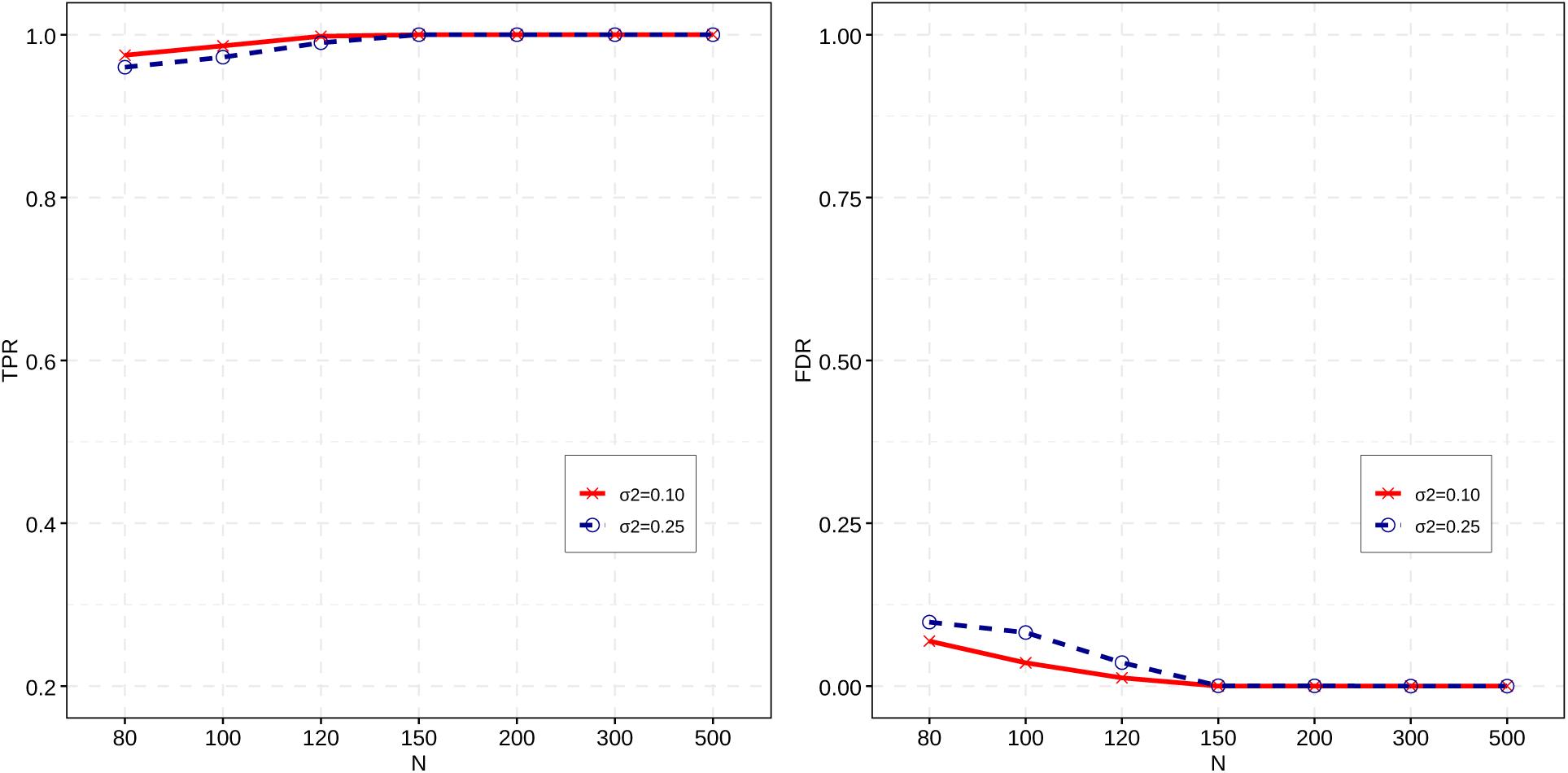
PD and FDR of detected network edges for the DAG networks. The number of sample *N* varies from 80 to 500 and noise variance *σ* ^2^ = 0.10 and 0.25. PD and FDR were obtained from 20 simulated network replicates.

**Figure S10.**
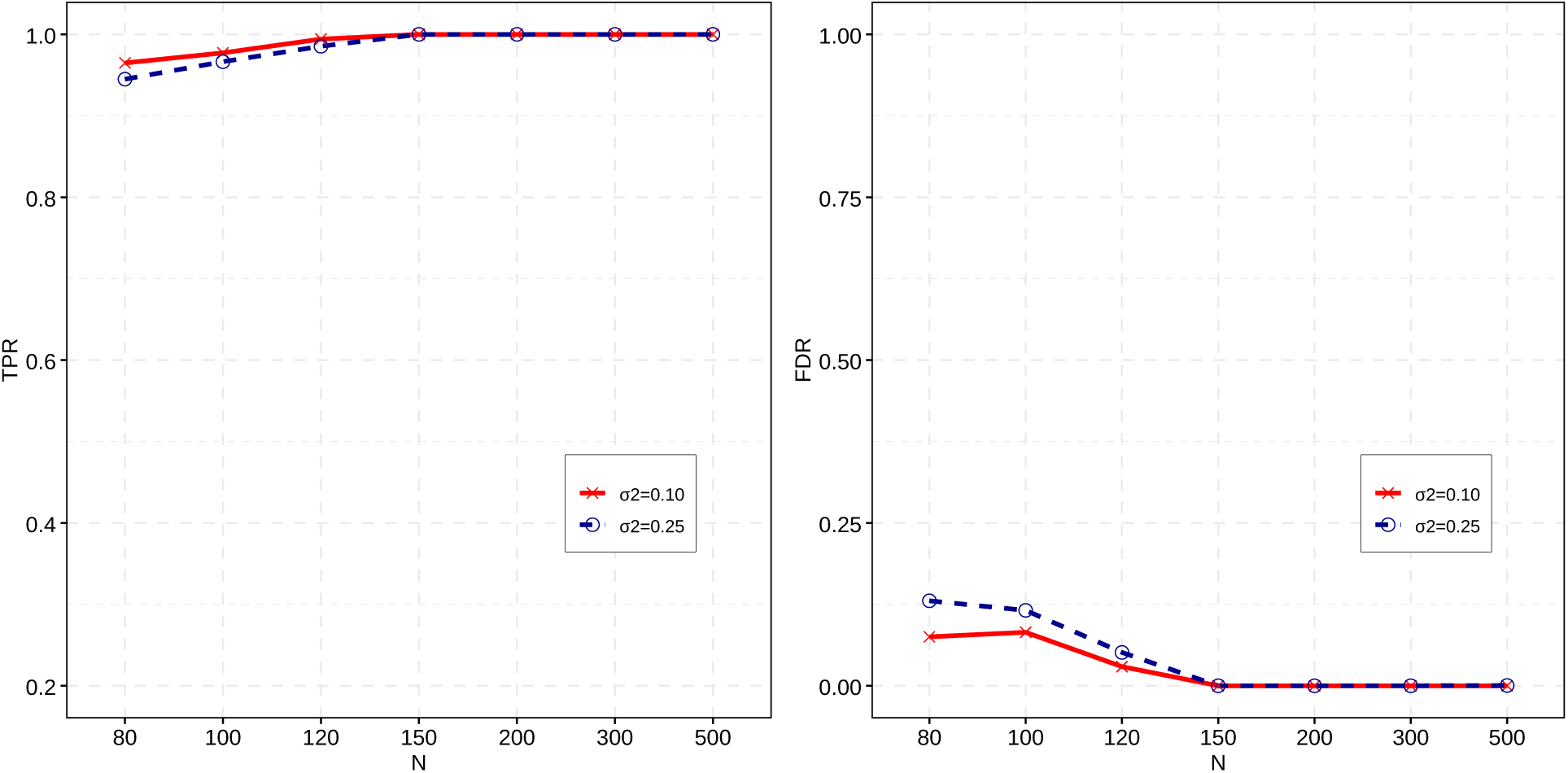
PD and FDR of detected network edges for the DCG networks. The number of sample *N* varies from 80 to 500 and noise variance *σ* ^2^ = 0.10 and 0.25. PD and FDR were obtained from 20 simulated network replicates.

